# Qoala-T: A supervised-learning tool for quality control of FreeSurfer segmented MRI data

**DOI:** 10.1101/278358

**Authors:** Eduard T. Klapwijk, Ferdi van de Kamp, Mara van der Meulen, Sabine Peters, Lara M. Wierenga

**Author notes:** **Corresponding author**: Eduard T. Klapwijk, Institute of Psychology, Leiden University, The Netherlands, Wassenaarseweg 52 2333 AK Leiden, The Netherlands., E.

## Abstract

Performing quality control to detect image artifacts and data-processing errors is crucial in structural magnetic resonance imaging, especially in developmental studies. Currently, many studies rely on visual inspection by trained raters for quality control. The subjectivity of these manual procedures lessens comparability between studies, and with growing study sizes quality control is increasingly time consuming. In addition, both inter-rater as well as intra-rater variability of manual quality control is high and may lead to inclusion of poor quality scans and exclusion of scans of usable quality. In the current study we present the Qoala-T tool, which is an easy and free to use supervised-learning model to reduce rater bias and misclassification in manual quality control procedures using FreeSurfer-processed scans. First, we manually rated quality of N = 784 FreeSurfer-processed T1-weighted scans acquired in three different waves in a longitudinal study. Different supervised-learning models were then compared to predict manual quality ratings using FreeSurfer segmented output data. Results show that the Qoala-T tool using random forests is able to predict scan quality with both high sensitivity and specificity (mean area under the curve (AUC) = 0.98). In addition, the Qoala-T tool was also able to adequately predict the quality of two novel unseen datasets (total N = 872). Finally, analyses of age effects showed that younger participants were more likely to have lower scan quality, underlining that scan quality might confound findings attributed to age effects. These outcomes indicate that this procedure could further help to reduce variability related to manual quality control, thereby benefiting the comparability of data quality between studies.

## Introduction

Quality control (QC) of structural magnetic resonance imaging (sMRI) data is an essential step in the image processing stream (Ducharme et al., 2016). These QC procedures enable detection of scanner-related artifacts, data-processing errors and participant-related artifacts such as head motion. Such artifacts are shown to result in underestimations of cortical volume and thickness measures (Reuter et al., 2015). This is of particular relevance to studies including children and adolescents or clinical groups (Brown et al., 2010; Van Dijk et al., 2012), as systematic group- and age-related differences in image quality (e.g., motion artifacts) may confound developmental or clinical inferences (Ducharme et al., 2016; Pardoe et al., 2016). Although QC procedures are widely applied, they are not extensively described in individual studies. Most studies use in-house developed protocols that include visual inspection of raw T1-weighted scans or output from automatic brain segmentation procedures (such as FreeSurfer; Dale et al., 1999). Trained raters then decide which scans are of sufficient quality for further analysis (see Backhausen et al., 2016 for a detailed example). The reliability of these protocols is currently unknown as different protocols are rarely compared and examined between studies. In general, manual quality ratings suffer from a number of limitations: they are inherently subjective, time consuming and susceptible to both inter-rater as well as intra-rater variability (e.g., learning or fatigue). Furthermore, not all artifacts might be detectable by visual inspection (Gardner et al., 1995). This together makes manual ratings vulnerable to 1) misclassification, resulting in either data loss or inclusion of poor quality data, and 2) inconsistency in thresholds that are used to determine which scans should be included (both within and between studies). These issues are also problematic for the replicability of studies as it complicates comparability of datasets (Poldrack et al., 2017). Hence there is a great need for automatized and unbiased QC assessments.

The present study introduces Qoala-T: A data-driven approach using a supervised-learning model to reduce rater bias and misclassification in manual QC procedures using FreeSurfer-processed scans. Our aim was to test which supervised-learning model would best distinguish good quality FreeSurfer-processed scans from scans of insufficient quality by comparing three often used supervised-learning algorithms (see Methods for details). Although several studies have described automatic tools to assess the quality of (raw) T1-weighted MRI data (Alfaro-Almagro et al., 2018; Esteban et al., 2017; Pizarro et al., 2016; White et al., 2018; Woodard & Carley-Spencer, 2006) these procedures did not focus on automatic segmented data (e.g. as processed in FreeSurfer) but were performed early in the processing stream. This is in contrast to manual QC procedures that are commonly assessed on segmented scans. We propose that a data-driven supervised-learning model in combination with manual QC on a subset of the FreeSurfer-processed data could overcome many of the issues related to manual QC. The procedure introduced in the present paper moves beyond earlier studies on automated quality assessment for a number of reasons: 1) We are one of the first to apply automated QC procedures on FreeSurfer processed data in a developmental dataset (see also Rosen et al., 2017 for QC of T1-weighted and segmented scans in a developmental sample; White et al., 2018 for QC of T1-weighted scans in a developmental sample). Developmental samples would benefit particularly from a thorough QC, given their increased susceptibility to artifacts including motion (e.g., Power et al., 2012). 2) The input measures for the supervised-learning model consisted of automatic segmented data (i.e., processed by FreeSurfer) such that it matches data used in our manual QC procedure. These processed data measures were fed into the supervised-learning models. The rationale for using processed data measures is that these measures are our data of interest and hence it is their quality that is the most crucial for further analysis. 3) Aside from within-sample cross-validation, we also applied the model on different novel independent datasets to test how well the model generalizes to other datasets. Hereby, we could demonstrate the potential of our current data-driven procedure, as our QC procedure could then be easily implemented elsewhere and extend to an unseen novel set of data. 4) To improve reproducibility we share a detailed description of our manual QC procedure that can be adapted to the specific needs of other datasets.

We propose that a full QC procedure using the Qoala-T tool would require manual QC only on a subset of data, either by retrospectively checking a subset of scans with indefinite quality scores or by using a rated subset as input for the supervised-learning model. Such a combination of manual QC and automated QC may be a sufficient and accurate way of assessing the quality of sMRI scans in other datasets as well. In addition, we provide R code of all steps used in the development of the Qoala-T tool in order to advance the reproducibility of the current analysis. Taken together, the automated QC procedure proposed in this study could critically increase comparability of QC procedures and as an added benefit also reduces the time needed for QC. This is particularly relevant with the increasing size of datasets and heterogeneity of multi-site labs.

## Materials and Methods

We used 784 T1-weighted scans acquired in three different waves from the BrainTime dataset, which is a large accelerated longitudinal research project normative development (Achterberg et al., 2016; Braams et al., 2015; Peters & Crone, 2017; Peters et al., 2016). For model evaluation, we also used 112 T1-weighted scans from an independent cross-sectional dataset that includes data from typically developing adolescents, adolescents with conduct disorder, and with autism spectrum disorder (BESD; Aghajani et al., 2016; Aghajani et al., 2017; Klapwijk et al., 2016a; Klapwijk et al., 2017; Klapwijk et al., 2016b), and 773 T1-weighted scans from 13 developmental sites from the Autism Brain Imaging Data Exchange dataset (ABIDE; http://fcon_1000.projects.nitrc.org/indi/abide/; Di Martino et al., 2014).

### Participants

The BrainTime study included a total of 784 T1-weighted scans collected over three time points, with approximately 2 year intervals (see Table 1). At the first time point, 299 participants between ages 8-25 years participated in this MRI study. A high-resolution structural T1-weighted scan was available for 292 participants (142 males). Mean age at time point one was 14.05 (SD=3.67; range=8.01-25.95). IQ was estimated at time point one with two subtests (Similarities and Block Design) from the WISC-III (for participants < 16 years) and the WAIS-III (for participants ≥ 16 years) (Wechsler, 1991, 1997). Mean IQ was 109.39 (SD=10.97, range=80-143). At the second time point two years later, T1-weighted scans were available for 252 participants (121 males). Mean age at time point two was 16.15 (SD=3.54, range=9.92-26.61). At time point two, IQ was estimated using two different subtests (Vocabulary and Picture Completion) from the WISC-III and the WAIS-III. Mean IQ at time point two was 108.24 (SD=10.26, range=80-148). At the third time point, T1-weighted scans were available for 240 participants (115 males). Mean age at time point three was 18.16 (SD=3.68, range=11.94-28.72). The grand total of included data was 784 T1-weighted scans. Exclusion criteria prior to participation were current use of psychotropic medication or a psychiatric diagnosis. Participants and their parents (for participants younger than 18 years old) provided written informed consent. Participants (<18 years) received presents for their participation and their parents received payment for travel costs. Participants (≥18 years) received payment for participation.

**Table 1.**
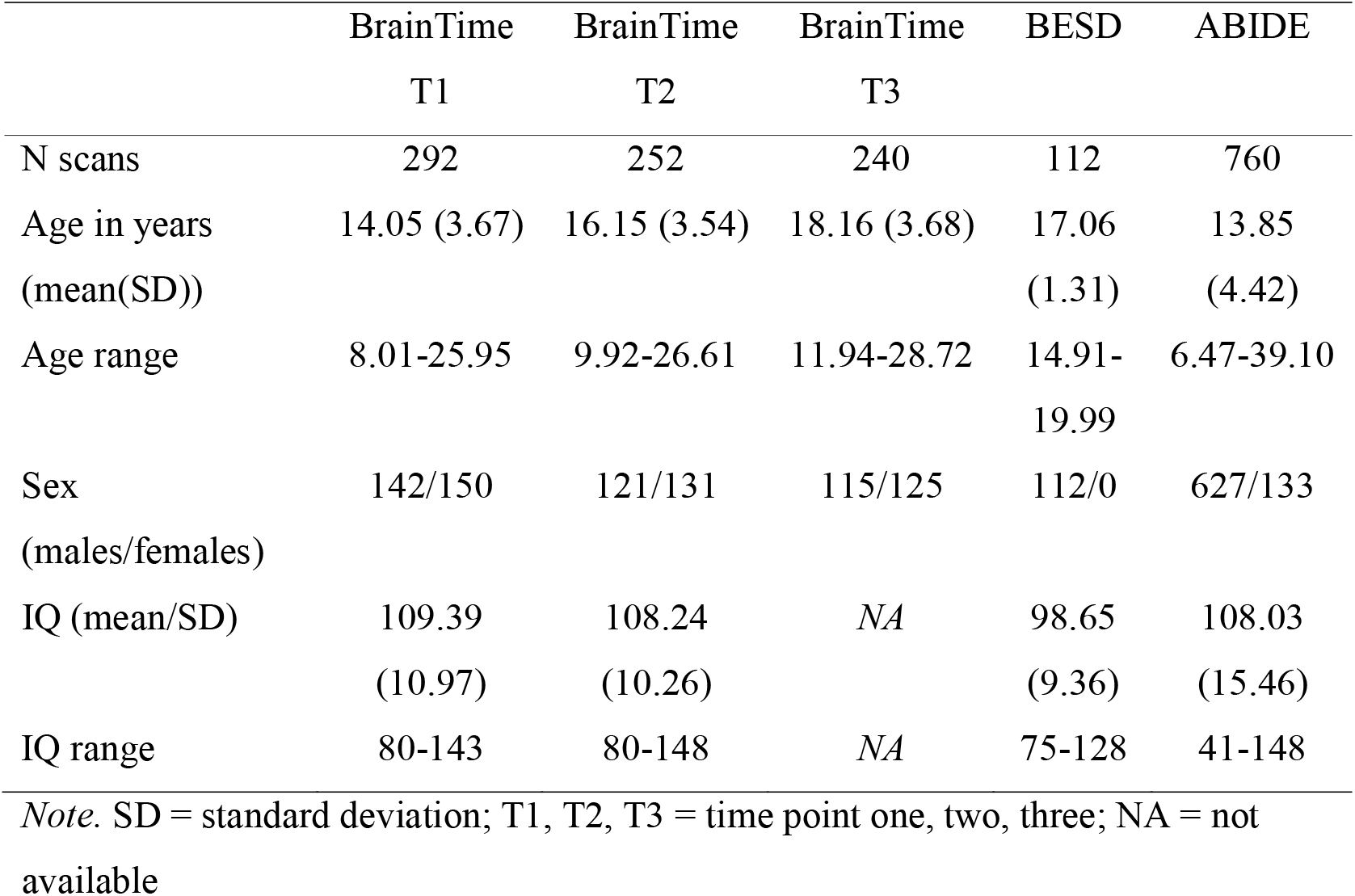
Demographic information for datasets

The BESD (Brain, Empathy, Social Decision-making) dataset consisted of 112 participants (all males). This included T1-weighted scans of 52 participants with conduct disorder, 23 participants with autism spectrum disorder, and 37 typically developing controls, 15-19 years old (mean 17.06, SD=1.31, range=14.91-19.99). IQ was estimated using the WISC-III (for participants < 16 years) and the WAIS-III (for participants ≥ 16 years) with Vocabulary and Block Design subtests. Mean IQ was 98.65 (SD=9.36, range=75-128). Exclusion criteria for participation in this study were neurological abnormalities, a history of epilepsy or seizures, head trauma, left-handedness, and IQ lower than 75. For more details regarding recruitment and clinical characteristics see Aghajani et al. (2017) and Klapwijk et al. (2016a).

The BrainTime and BESD study were approved by the Leiden University Medical Ethical Committee. A radiologist reviewed all anatomical scans and no clinically relevant abnormalities were detected.

T1-weighted scans (N = 773; after 13 failed FreeSurfer processing final N = 760) from the following sites from the ABIDE dataset (Di Martino et al., 2014) were used: Kennedy Krieger Institute, NYU Langone Medical Center, Olin Institute of Living at Hartford Hospital, Oregon Health and Science University, San Diego State University, Stanford University, Trinity Centre for Health Sciences, University of California Los Angeles sample 1 and 2, University of Leuven sample 2, University of Michigan sample 1 and 2, and Yale Child Study Center.

### Imaging Data Acquisition and Processing

MRI scans of the BrainTime and BESD samples were acquired on a Philips Achieva TX 3.0 T scanner, while using a standard whole-head coil. To minimize head motion, participants were familiarized with the scanner environment using a mock scanner, their heads were fixated using foam pillows, and the importance of lying still was emphasized to participants in between scan sequences. A high-resolution 3D T1 anatomical scan was acquired a (TR = 9.76 ms, TE = 4.59 ms, 140 slices, voxel size = 0.875 × 0.875 × 1.2 mm, FOV= 224 × 177 × 168 mm). Acquisition parameters for the different ABIDE sites are online available at http://fcon_1000.projects.nitrc.org/indi/abide/.

Tissue classification and anatomical labeling was performed on the basis of the T1-weighted scan using the well-validated and well-documented FreeSurfer v6.0.0 software (http://surfer.nmr.mgh.harvard.edu/). In short, this software includes tools for non-brain tissue removal, cortical surface reconstruction, subcortical segmentation, cortical parcellation, and estimation of various measures of brain morphometry. Technical details of the automated reconstruction scheme are described elsewhere (Dale et al., 1999; Fischl et al., 2002; Fischl et al., 1999). The crosssectional FreeSurfer outputs were used for all datasets (including the longitudinal BrainTime sample). After preprocessing, measures of cortical thickness and surface area based on the Desikan-Killiany atlas (Desikan et al., 2006) and subcortical volumes (Fischl et al., 2002) were computed (i.e., aparc_thickness_lh/rh, aparc_area_lh/rh, aseg_stats files using stats2table command) and used as input measures for the supervised-learning models (see Supplementary Table 1 for full list of variables). Note that we did not include cortical volume measures, as cortical volume is a composite measure of cortical thickness and surface area. Anterior frontal and temporal poles were not included in the estimates of the outcome measures, due to frequent bad segmentation of these regions (see also Keshavan et al., 2018a).

### Manual Quality Control

In order to relate outcomes of the supervised-learning models to outcomes of the manual quality control, we first rated all FreeSurfer-processed scans from the BrainTime and BESD samples manually. Five raters (E.K., F.K., M.M., S.P., L.W.) with experience in similar QC protocols were trained to perform the manual QC procedures developed in our lab, see Supplementary Materials. The training set for the current protocol included 20 scans from an independent developmental dataset (not described). Next, the BrainTime dataset was randomized and split into five subsets. Each scan was rated as 1 = ‘Excellent’, 2 = ‘Good’, 3 = ‘Doubtful’, or 4 = ‘Failed’, based on a set of specific criteria (e.g., affection by movement, missing brain areas in reconstruction, inclusion of dura or skull in reconstruction). A detailed, step-by-step explanation of the manual QC procedure can be found in the Supplementary Materials. To test reliability between the five raters, each subset of the BrainTime dataset included 10% overlap (i.e., 80 scans) such that these 80 scans were rated 5 times. In order to control for intra-rater drift, such as practice effects, each rater assessed half (N=40) of the overlapping scans at the start of the QC procedure, and the other half of the overlapping scans at the end of the QC procedure. Intra-class correlations were computed as an indication of inter-rater reliability. For the ABIDE dataset, we used manual ratings released by the MRIQC project (https://github.com/poldracklab/mriqc; Esteban et al., 2017). These scans were rated on a three-point scale (accept/doubtful/exclude) by two different raters, aided by FreeSurfer surface reconstructions (see Esteban et al., 2017 for details). Distribution of manual QC labels for the different datasets is reported in the Supplementary Materials (see Supplementary Table 2).

### Supervised-learning: the Qoala-T tool

This paper introduces a supervised-learning tool (Qoala-T) to assess manual quality ratings of FreeSurfer automatic segmented T1-weighted scans. Different supervised-learning models were compared. For these predictions, we used binary classification of included and excluded scans: manual rating scores ‘Excellent’, ‘Good’, and ‘Doubtful’ were considered scans to be included and manual scores ‘Failed’ were considered exclusions. Our aims were to test 1) which supervised-learning model would best predict scan quality, 2) whether manual ratings of a subset of the data (e.g. as low as 10%) would be able to predict the scan quality of the other 90% of the data, 3) which cutoff for exclusion should be used (i.e., whether scans that were manually rated as ‘Doubtful’ should be excluded in addition to ‘Failed’ scans) and 4) performance of the model on novel datasets (BESD and ABIDE samples). Binary classification was performed because even when using multiple categories for manual ratings (e.g., four categories in our case), we eventually want to determine which scans should be excluded and which scans should not. All statistical analyses were performed in R version 3.3.3 (R Core Team 2014). The development of the Qoala-T tool included a number of steps as described below.

### Step 1: Model selection and hyper parameter tuning

First, we performed a preliminary evaluation to select the best performing supervised-learning model using 60% of the BrainTime data (see Figure 1A for model procedure).

**Figure 1:**
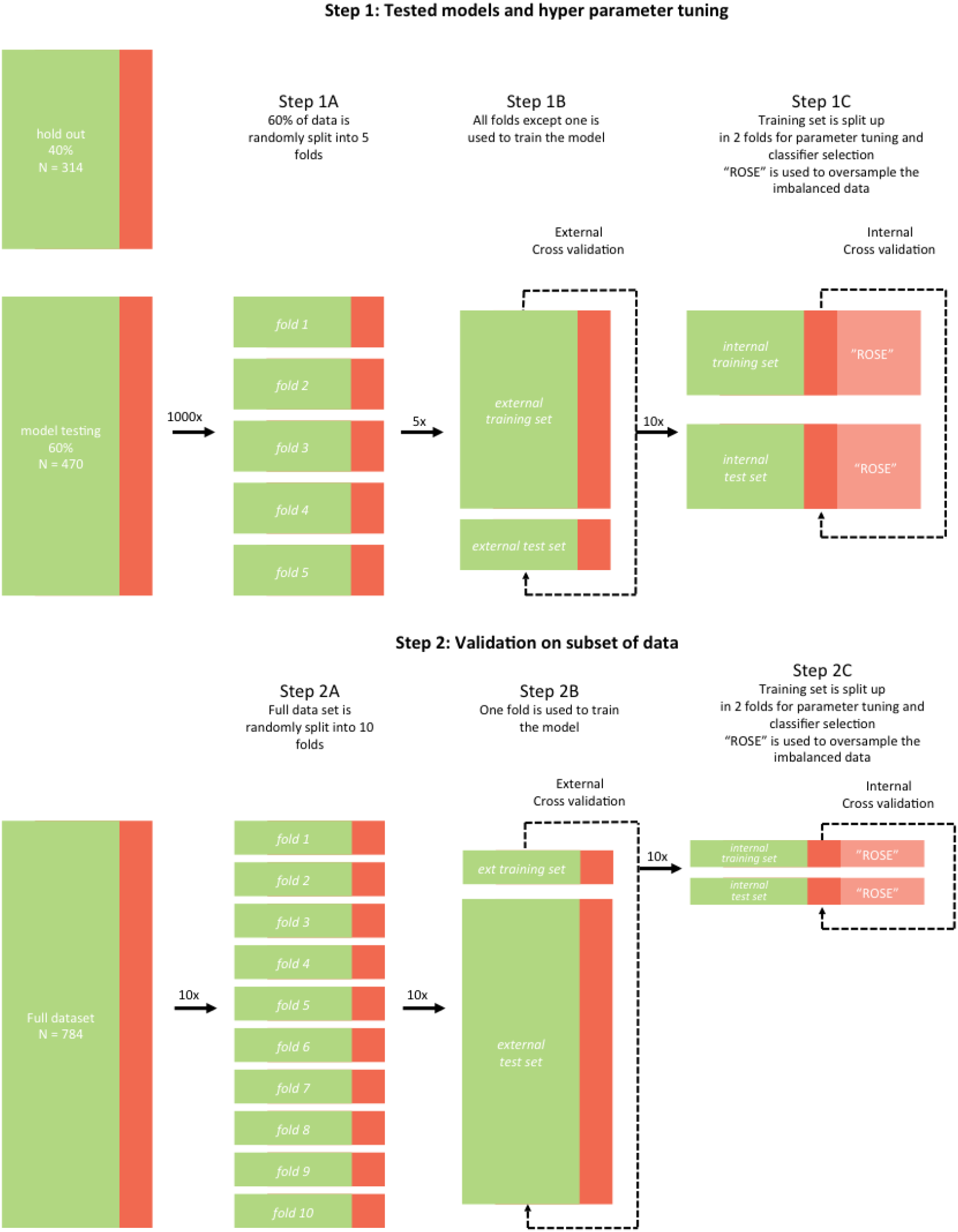
Illustration of Step 1 (Model selection and hyper parameter tuning) and Step 2 (Validation on subset of data) of development of the Qoala-T tool. Green indicates included scans based on manual ratings, red indicates scans manually rated to be excluded.

Two of the most often used supervised-learning models in predicting MRI quality are support vector machine (SVM) (Esteban et al., 2017; Pizarro et al., 2016) and random forests modeling (RF) (Esteban et al., 2017). For the purpose of this paper we compared these models in addition to another tree-based supervised-learning model (gradient boosting machines; GBM) using the caret R library (Kuhn, 2008). SVM is a model in which input vectors are mapped into a high-dimensional space in which different categories are optimally separated by a hyperplane; new samples are then classified based on which side of the hyperplane they fall (Cortes & Vapnik, 1995). In RF models, a number of decision trees are generated for bootstrapped samples of the data, in which features are compared against a threshold.

Classification of new data then occurs by popular vote over all trees (Breiman, 1996, 2001). GBM is also a tree-based statistical learning tool, but it differs from RF in that outcomes of previous trees influence subsequent trees by adjusting weights of misclassified instances (Freund & Schapire, 1995). These supervised-learning models each have a number of hyper-parameters that can be adjusted to optimize classification. To estimate the most optimal hyper-parameters, all models were tested using different combinations of hyper-parameter settings. Ultimately, we evaluated the best performing model using the most optimal hyper-parameter settings for that algorithm.

The analysis pipeline for this step is shown in Figure 1. First, the data was partitioned into a hold out and model selection dataset. Next, for each supervised-learning model, the model selection data set was divided into five folds. As such, 5fold nested cross-validation was repeated 1000 times (external cross validation, step 1A and 1B in Figure 1). Within each of these 5 folds, again 2-fold cross-validation was repeated 10 times for classifier selection and hyper-parameter tuning (internal cross validation). For each of the 1000 repetitions results were averaged over the 5 folds resulting in 1000 AUC estimates per supervised-learning model. The three models were compared using ANOVA and post-hoc t-tests. The hyper-parameters leading to the maximum average of the AUC scores in step 1C (internal cross validation) were selected for further analysis. To overcome issues related to imbalanced classes (a lower number of excluded scans than included scans: in the BrainTime sample 48 out of 748 were manually rated ‘Failed’) we used ‘Random Over-Sampling Examples’ (ROSE; Lunardon et al., 2014; Menardi & Torelli, 2014). ROSE is a bootstrap technique that generates synthetic balanced samples to improve estimations of binary classifiers that can be implemented within ‘caret’.

### Step 2: Validation on subset of data

In the second step, we used the best supervised-learning model (SVM, RF or GBM) and the best hyper-parameter setting for that model selected in step 1, to evaluate the validity of the model. The aim of this step was to assess whether manual QC on only a subset of the scans (as low as 10%) would be sufficient to predict which scans should be excluded from the total sample. To do so we included 10% of the BrainTime data as the external cross validation training set and 90% of the dataset as the external test set, this step was repeated 10 times. Within these folds, 10 repeats of 2-fold internal cross validation was performed for parameter tuning and classifier selection (see Figure 1B).

### Class probability as an indicator of scan quality

Predicted outcome (inclusion or exclusion) and class probabilities were estimated for each scan. Class probabilities were used as an estimate of scan quality where an estimated scan quality above 50% indicated good quality whereas a probability lower than 50% indicated poor quality. The binary prediction model predicts ‘exclude’ when scans have poor quality below 50% and predicts ‘include’ for scans with good quality above 50%. As such, scan quality estimates closer to 50% may indicate higher uncertainty of the model to predict the binary outcome. To test the stability of the class probability estimates, validation was repeated 10 times, resulting in 10 folds (10-fold external cross-validation), such that all scans were part of the training set exactly once and part of the testing set 9 times. Within each fold output values (AUC, sensitivity, specificity and feature ranking) were assessed using 10 repetitions of 2fold internal cross-validation (see Figure 1B). For each scan predicted classification and class probability estimates were used to test goodness of model fit.

### Step 3: Where to draw the line – evaluation of QC threshold

In this step we evaluated the threshold for exclusion of scans. In previous steps scans manually rated as ‘Failed’ were excluded. However, perhaps a more stringent threshold where scans rated as ‘Doubtful’ are also labeled as ‘exclude’ would be a more sufficient cutoff. We repeated step 2 but this time labeled scans rated both as ‘Doubtful’ and ‘Failed’ as eligible for exclusion. The mean AUC values predicted by the 10-fold cross validation were compared to the AUC values of the threshold model described above, using a one sided t-test.

### Step 4: Evaluation on novel datasets

The model based on BrainTime data was tested on the BESD data (N = 112; see Participants section for details), of which all FreeSurfer-processed scans were manually rated. Secondly we assessed whether manual QC on only a subset of BESD data (10%) would be able to predict which scans should be excluded from the total sample. For this, we used 10-fold cross validation in which in each fold 10% of the scans were used as the training set (see step 2). As such, each scan was used in the training set once, and for each scan the outcome (include/exclude) was predicted 9 times. In addition, the model based on BrainTime was also tested on ABIDE data (N = 760; see Participants section for details), using manual ratings released by the MRIQC project (https://github.com/poldracklab/mriqc; Esteban et al., 2017). Note that the MRIQC manual rating procedure was different from the procedure used for the BrainTime model, allowing us to compare Qoala-T scores with a different manual rating procedure rated by different raters. This is important, since raters and procedures will also vary for future datasets that use Qoala-T.

### Additional analyses: relation with age, sex, and MRIQC metrics

To investigate age and sex effects on scan quality, we explored the relations between Qoala-T scores and age and sex using linear regression in the BrainTime, BESD and ABIDE datasets separately (in the BESD dataset only age effects were explored, as this sample is male only). We also explored the relation between Qoala-T scores and MRIQC, an existing automatic QC procedure that evaluates raw imaging data (Esteban et al., 2017). All T1-weighted scans in the BrainTime and BESD datasets were processed using MRIQC version 0.11.0, for ABIDE the MRIQC metrics (processed using MRIQC version 0.9.6) available online (https://github.com/poldracklab/mriqc/blob/master/mriqc/data/csv/x_abide.csv) were used (see Supplementary Table 3 for full list of MRIQC metrics). Relations between Qoala-T scores and MRIQC metrics were assessed using multiple linear regression in the BrainTime, BESD and ABIDE datasets separately.

## Results

### Correspondence of manual QC amongst raters

To test rater correspondence, 80 scans (~10%) of the BrainTime dataset were rated by all five raters. Even though all raters followed the same protocol and were trained extensively, only 7.5% of the scans (i.e. 6 out of 80 scans) were rated in complete agreement. Of the 80 scans in this set, 86.3% of the scans (69 scans) were rated with an agreement of at least three raters (i.e. at least three raters gave the same score). For these scans, the most frequent rating was used as the final score. For the remaining scans with no majority rating (e.g., rated twice as Excellent, twice as Good, and once as Doubtful), the rating of the rater with the overall highest reliability (i.e., the rater with highest mean intraclass correlation coefficient (ICC) reliability in pairwise comparisons with the other raters) was used. Note that in these cases there was always one other rater who had given the same rating as the most reliable rater. The intercoder reliability for the quality of the scans ranged from .38 to .72, with a mean intercoder reliability of .53 (all *p* < .001), indicating moderate reliability (Portney & Watkins, 2000). When using two categories as in the binary prediction model (include/exclude), the mean inter-coder reliability was comparable (.49), ranging from .25 to .79. The results of the five ratings per scan can be observed in Figure 2. This figure illustrates the subjective aspect of manual QC assessment, even amongst well-trained raters, herewith highlighting the need for objective assessment.

**Figure 2:**
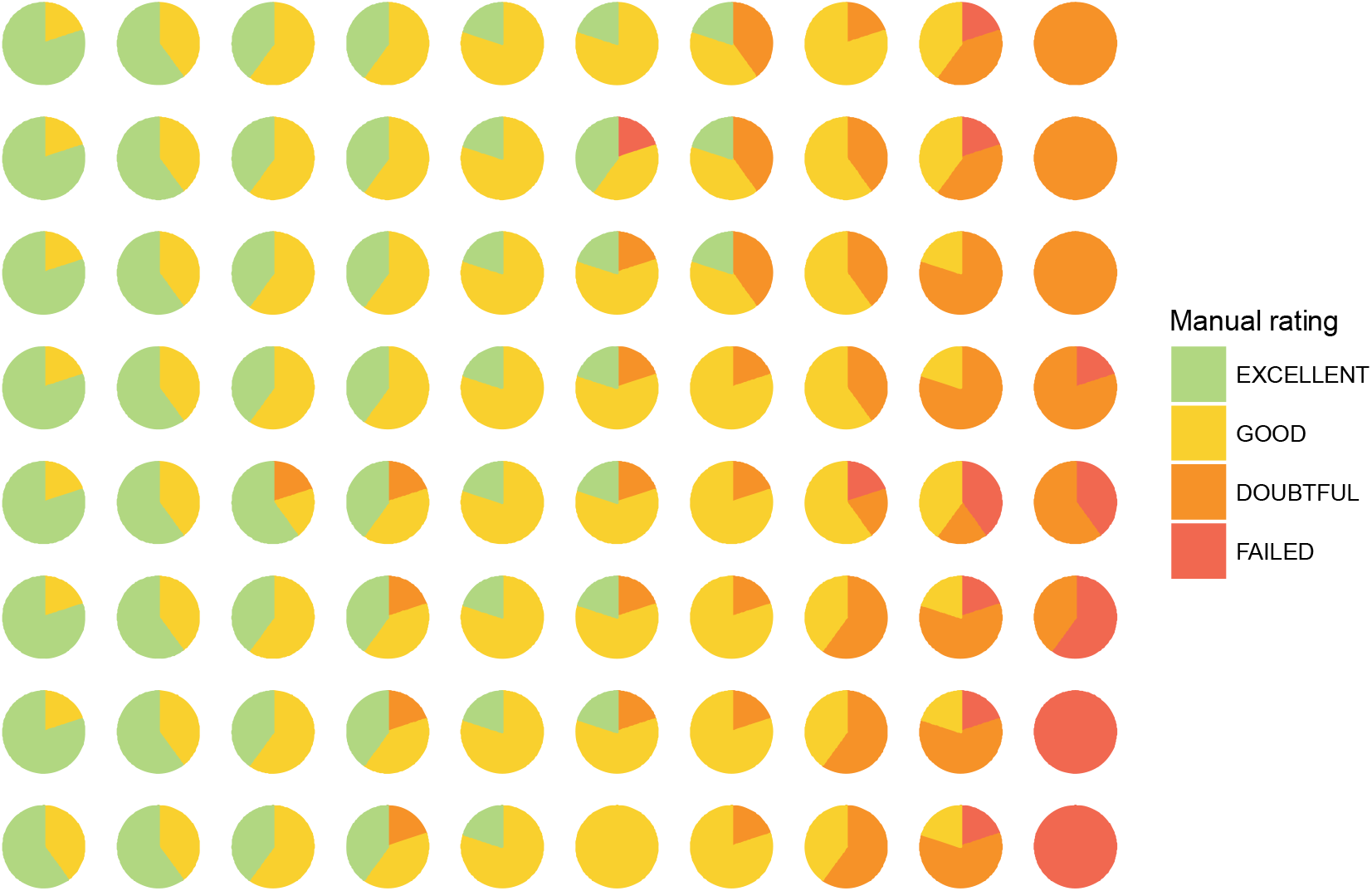
A subset of 80 scans was subjected to manual QC by all five raters. Each pie represents one scan and each pie slice represents a rater where the color of that piece indicates the final rating given by a single rater (e.g. Excellent, Good, Doubtful, Failed).

### Model selection and hyper parameter tuning

In step one the performance of the three supervised-learning models (SVM, RF, and GBM) was compared to test which of the three models was the most accurate in predicting manual quality ratings. The best supervised-learning model was selected based on the AUC values. The AUC results of the model comparison can be observed in Figure 3. Analysis of variance showed that performance differed across the three algorithms (F(2, 2909) = 1270, p < .001), see results in Table 2 and Figure 3. A post hoc t-test showed that the AUC of the RF model was higher compared to both the GBM model (p < .001) as well as the SVM model (p < .001). Hence, the RF model was used in the next set of analyses, where the best performing hyper-parameter settings were 501 trees to grow (i.e., *ntree*) and 8 variables randomly sampled as candidates at each split (i.e., *mtry*).

**Table 2.** Model comparison of supervised-learning models to predict manual qualiy ratings of FreeSurfer-processed T1-weighted BrainTime scans

|  | AUC |  | Sensitivity |  | Specificity |  |
| --- | --- | --- | --- | --- | --- | --- |
|  | mean | SD | mean | SD | mean | SD |
| RF (random forests) | 0.921 | 0.013 | 0.806 | 0.042 | 0.859 | 0.083 |
| GBM (gradient<br>boosting machine) | 0.867 | 0.040 | 1.000 | 0.000 | 0.068 | 0.001 |
| SVM (support vector<br>machine) | 0.880 | 0.005 | 0.573 | 0.014 | 0.944 | 0.002 |

*Note.* AUC = area under the curve; SD = standard deviation.

**Figure 3.**
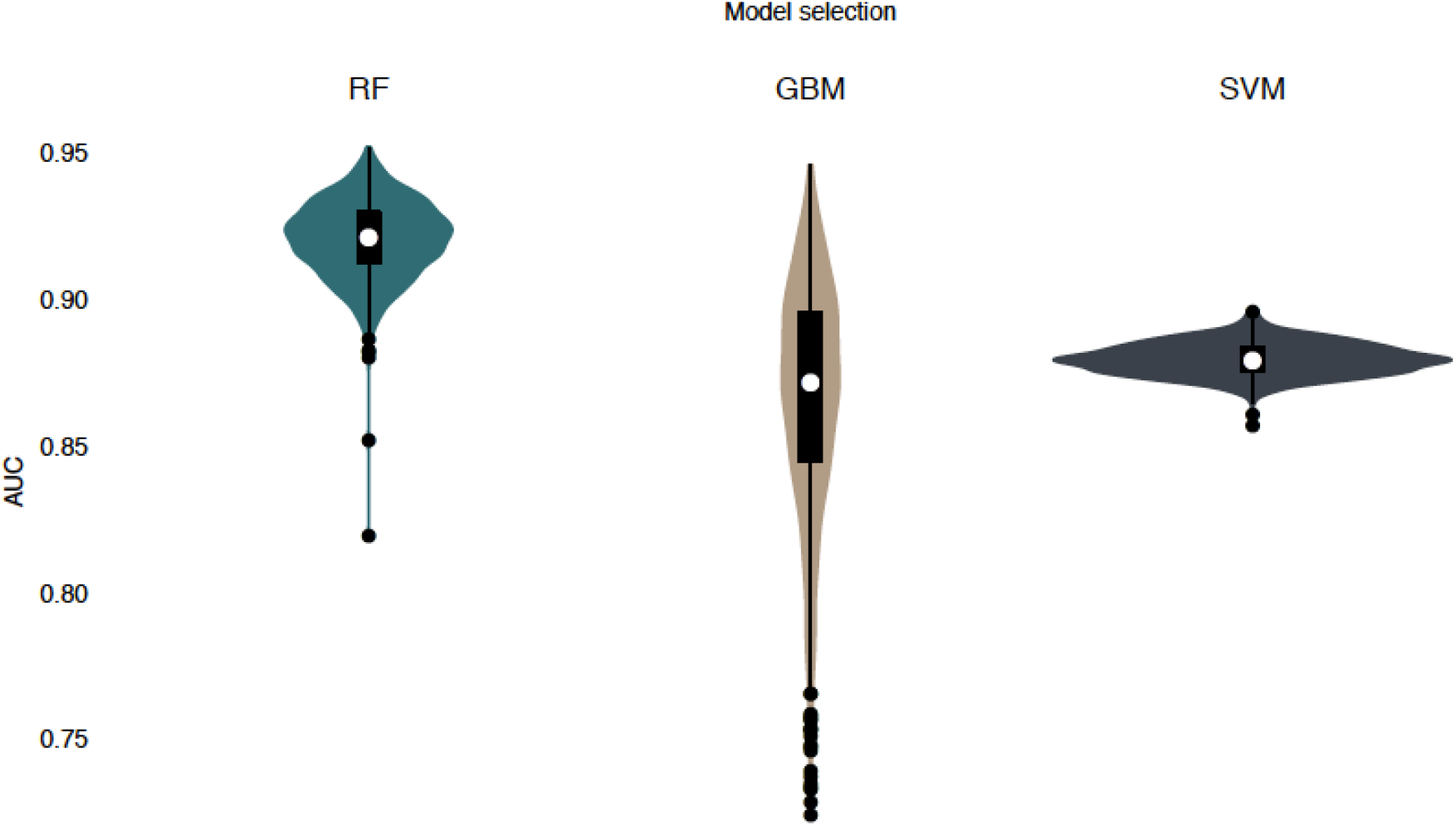
Violin plot of area under the curve (AUC) values 1000 repetitions averaged over 5 folds for random forests (RF), gradient boosting machine (GBM) and support vector machine (SVM) models. The graphs show the spread of the AUC values for all folds and repetitions. Higher AUC values indicate better model fit. For instance, for GBM, there is a large spread in how well the model performed across all folds and repetitions, whereas for the RFmodel the mean AUC was higher and also showed less dispersion.

### Validation using subset of data

The next set of analyses assessed how well the model was able to predict quality rating when only a subset of the sample (in our case 10%) would have had manual QC. To test the stability of the results this step was repeated 10 times. To do so we used 10-fold cross validation where in each fold 10% of the scans were used as the training set, in this way each scan was used in the training set once, and for each scan the outcome (include/exclude) was predicted 9 times. To estimate model accuracy these predictions were compared to the manual rating. The results of the 10 folds can be observed in Table 3. Mean AUC of the 10-folds was .977 (SD = .005). The results show that with 10 % of the data being manually assessed, whether a scan should be included or excluded in the remaining 90 % of the data quality can be predicted with both high sensitivity and specificity. The model provides not only a binary outcome (include/exclude) ‘ but also a probability score which indicates with how much certainty that scan was classified in the included or excluded group. This measure was used as an indication of scan quality (ranging from 0 to 100). Potentially these probability levels could be used to flag scans that would benefit from manual QC in a novel dataset. Because each scan was predicted by the model nine times, there were nine of these inclusion/exclusion probabilities estimated for each scan see Figure 4. These nine inclusion/exclusion probabilities were averaged to assess the predicted exclusion rate. As can be seen in Figure 4, scans with extremely high or extremely low scan quality estimates are more likely to be classified correctly than scans with average probability. In addition, scan quality ratings at the extreme ends showed more stable predictions across the 10 folds. The total number of falsely excluded scans was 21 out of 738 included scans (2.85 %) and the number of false negatives was 5 out of 46 excluded scans (10.9 %) see grey dots in Figure 4.

**Table 3.** 10-fold cross validation using subsets of BrainTime data to predict manual quality ratings of FreeSurfer-processed T1-weighted BrainTime scans

| Fold | AUC | Accuracy | Sensitivity | Specificity | PPV | NPV |
| --- | --- | --- | --- | --- | --- | --- |
| 1 | 0.973 | 0.966 | 0.902 | 0.970 | 0.649 | 0.994 |
| 2 | 0.971 | 0.963 | 0.878 | 0.968 | 0.632 | 0.992 |
| 3 | 0.984 | 0.969 | 0.881 | 0.974 | 0.685 | 0.992 |
| 4 | 0.977 | 0.969 | 0.829 | 0.977 | 0.694 | 0.989 |
| 5 | 0.969 | 0.962 | 0.905 | 0.965 | 0.623 | 0.994 |
| 6 | 0.979 | 0.962 | 0.878 | 0.967 | 0.621 | 0.992 |
| 7 | 0.980 | 0.967 | 0.881 | 0.973 | 0.673 | 0.992 |
| 8 | 0.976 | 0.959 | 0.881 | 0.964 | 0.607 | 0.992 |
| 9 | 0.979 | 0.963 | 0.902 | 0.967 | 0.627 | 0.994 |
| 10 | 0.980 | 0.965 | 0.829 | 0.973 | 0.654 | 0.989 |
| Mean | 0.977 | 0.964 | 0.877 | 0.970 | 0.646 | 0.992 |
| SD | 0.005 | 0.003 | 0.027 | 0.004 | 0.030 | 0.002 |

*Note.* AUC = area under the curve; PPV = positive predictive value (i.e., proportion of scans tested positive as exclusions that were correctly predicted to be excluded); NPV = negative predictive value (i.e., proportion of scans tested negative as exclusions that were correctly predicted to be included); SD = standard deviation.

**Figure 4.**
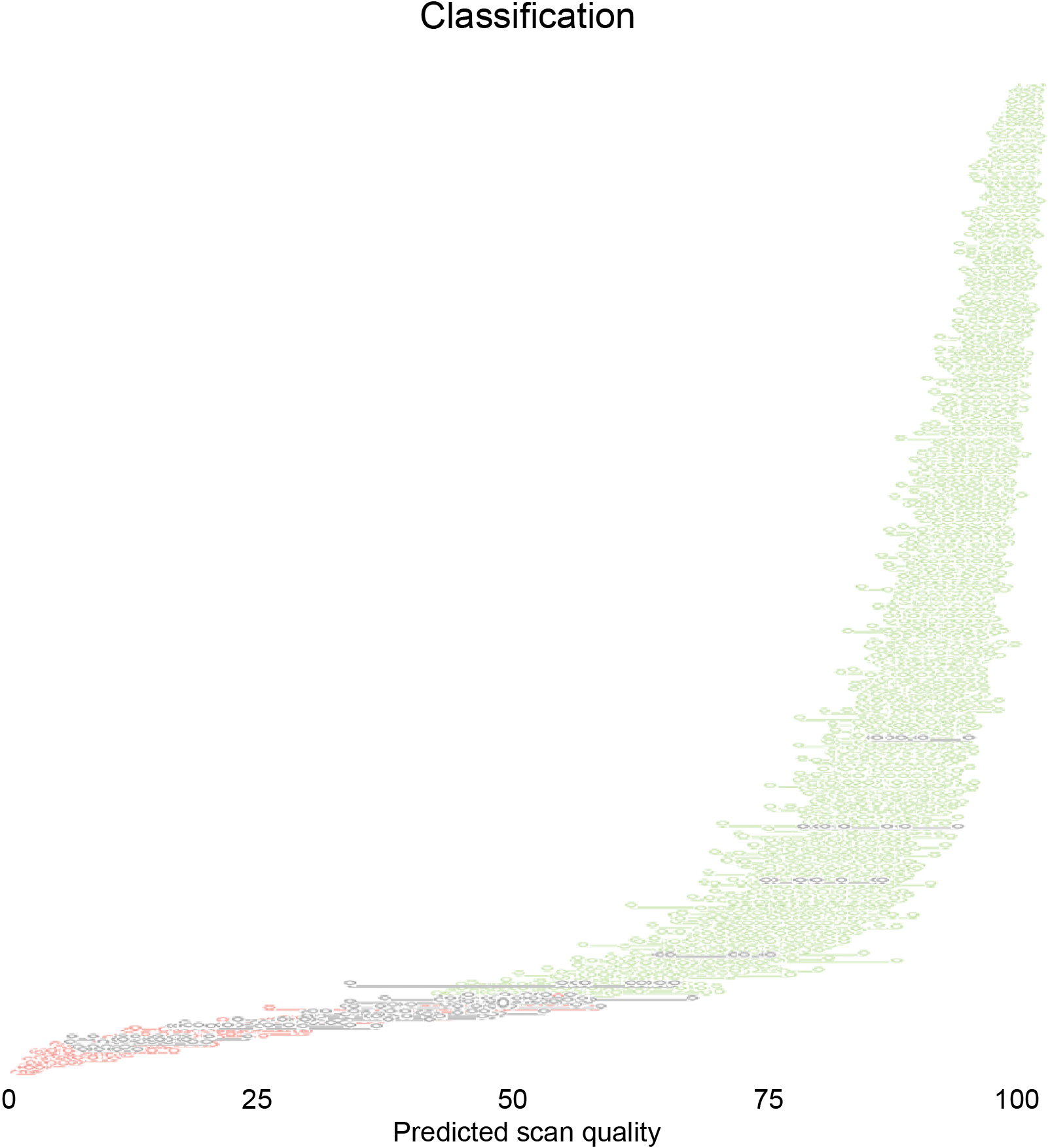
For each scan 9 values for predicted scan quality (0-100) are displayed on the x-axis. Each value on the y-axis represents a scan ordered by mean scan quality. Lines between dots connect 9 values of predicted scan quality for a single scan. Scans with an average scan quality <50% are predicted to be excluded, and scan with an average scan quality =50% are predicted to be included. Colors indicate goodness of predicted classification green = ‘correct include’ red = ‘correct exclude’ grey = ‘incorrect classification’). A total of 26 scans out of a total of 784 scans (3.3%) were misclassified.

It could be the case that the AUC is simply high due to the imbalanced exclusion rate. That is, because of the low number of exclusions, it could be that the best model would simply predict to include all scans, which would result in relatively high levels of model prediction accuracy. In order to check the above chance model accuracy, we reassessed AUC by randomly shuffling exclusion labels (1000 permutations). AUC values of our model were considerably higher than the permuted AUC values (mean AUC = .502, SD = .052, p < .001), meaning that the model performed much better than a model in which scans were randomly classified. In addition, the relatively low variance of the results of the validation scores (see Table 3) suggests that ROSE served as an adequate oversampling method (Lunardon et al., 2014).

### Class probabilities as indicator of scan quality

We compared mean class probabilities of the different manual rating categories for scans that were predicted to be included and excluded by the model, since this indicates with how much certainty the model was able to predict the binary outcome (include/exclude). A total of 717 scans were rated as ‘include’ by both manual ratings and the model (correct inclusions), and 41 scans were rated as ‘exclude’ by both the manual ratings and the model (correct exclusions). There were 21 scans that were excluded by the model but not by manual ratings (false excluded scans), and 5 scans that were included by the model but excluded according to manual ratings (false included scans). These incorrectly classified scans could be related to errors in model prediction, or errors in manual rating (e.g., due to fatigue). Hence, further investigation of scan quality levels (i.e. with how much certainty the model places a scan in the ‘include’ or ‘exclude’ category) could provide insights into the goodness of model fit. The mean class probability levels for each scan estimated in step 2 are displayed in Figure 5, in which the manual rating score is color-coded. Interestingly, 17 of the 21 ‘false’ excluded scans were already manually rated as ‘Doubtful’, and it could therefore be that they are in fact scans that should be excluded (i.e., potentially an error in manual ratings). Most scans manually rated as Doubtful (n = 143) were correctly identified as scans to be included. In addition, the mean probability scores of scans for which the manual and model rating differed were lower than when these were in concordance, showing that the model was already less confident about these ratings. That is, the mean scan quality score of false inclusions (red) was lower than the mean scan quality of true inclusions (orange, yellow and green). Similarly, false excluded scans (orange, yellow and green) have a higher mean scan quality compared to correct excluded scans (red). Note that these 4-category manual ratings were not included in model building, as only binary classification (include/exclude with cutoff between 3 ‘Doubtful’ and 4 ‘Failed’) was estimated.

**Figure 5.**
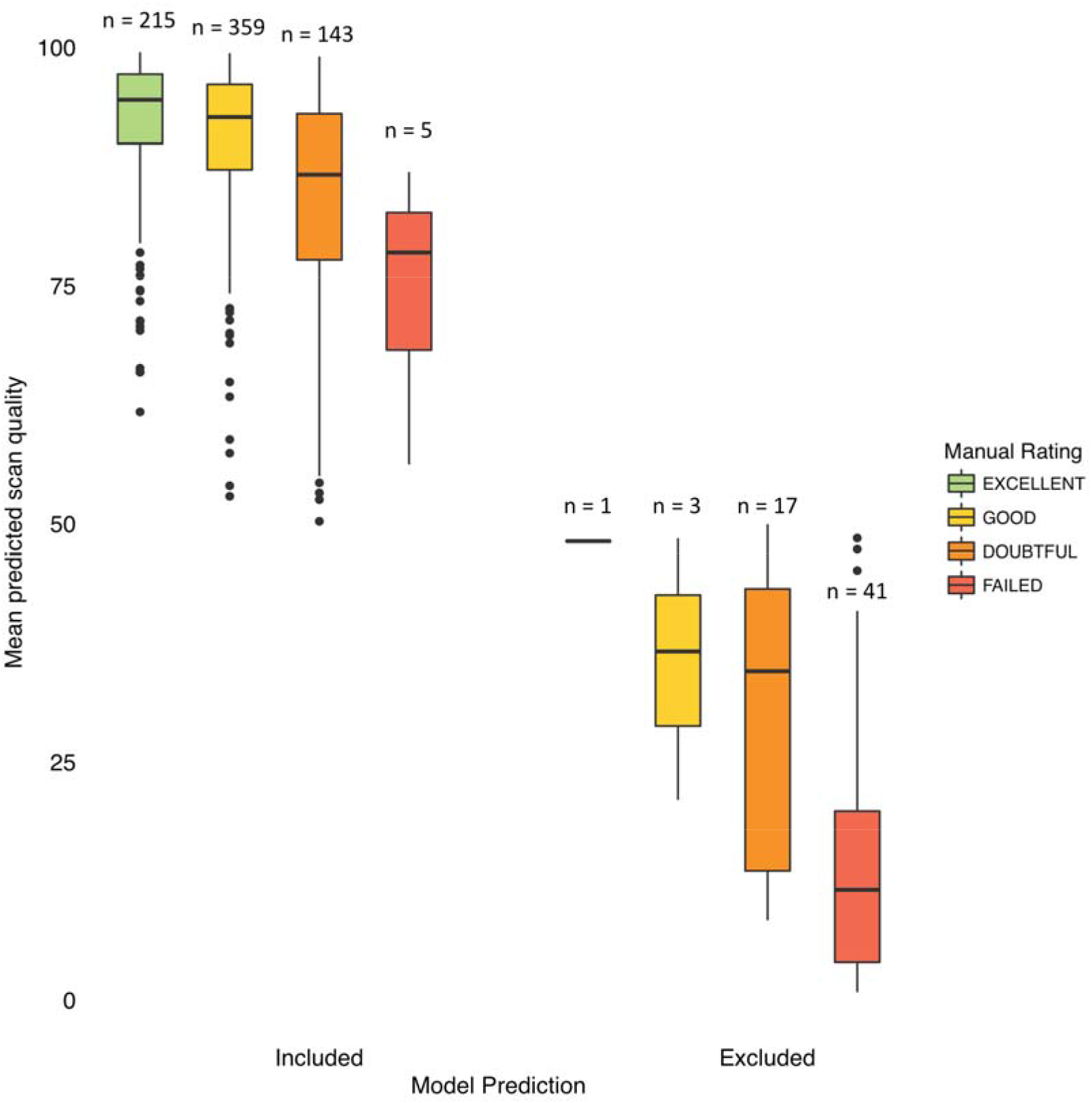
Boxplots showing mean scan quality probabilities as predicted by the Qoala-T model, grouped by binary model predictions (included/excluded) and subsequently by manual rating category. Colors of the boxplots indicate manual rating categories green = ‘Excellent’ yellow = ‘Good’, orange = ‘Doubtful’, red = ‘Failed’). Scans with a scan quality level < 50% are predicted to be excluded (right), and vice versa scans with scan quality levels > 50% are predicted to be included (left).

### Variable importance for model prediction

Which variables are most important for the prediction of quality rating? The random forests model provides additional information about the importance of explanatory variables using marginal permutation. This method randomly permutes an explanatory variable (e.g. left caudate thickness) and shows the mean decrease in prediction error that results from this permutation. If the explanatory variable is important for the accurate prediction of the model the predictive power of the model should systematically *decrease* model performance, while an unimportant variable would leave the predictive value unaffected (Breiman, 2001; Strobl et al., 2008). Variables with the top 50 highest importancy values for the BrainTime sample are summed in Figure 6 (for visualization purpose only the top 50 out of 185 variables are shown in the Figure, see Supplementary Table 1 for the full list of 185 variables and their importancy values). Mean importance variables across 10-fold cross validation on the full BrainTime sample showed highest importance values for left and right surface holes followed by white matter hypointensities, right entorhinal, and left and right precentral thickness.

**Figure 6.**
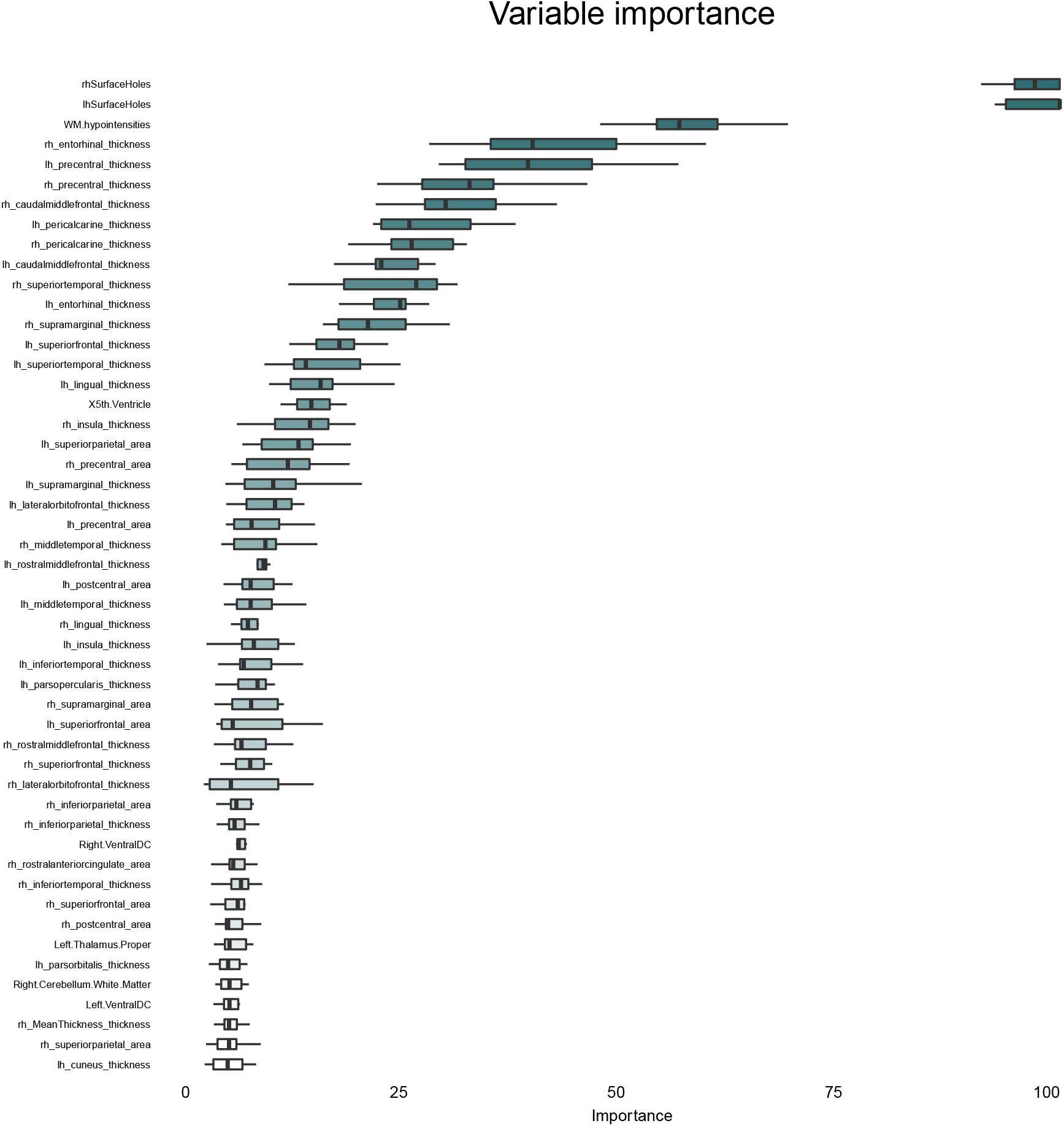
The permutation importance of explanatory variables for the prediction of scan quality (all 185 variables were used for the prediction but only the top 50 variables with the highest importance values are displayed). The importance measure on the x-axis represents the mean increase in classification error that results from randomly permuting that variable. Each boxplot shows the importance values of the 10 folds, ordered by their mean. Note that the importance value is a relative value that allows for a quick assessment of the relevance of a predictor for the outcome of interest. See Supplementary Table 1 for a full list of variables.

### Where to draw the line – evaluation of manual QC threshold

We also investigated whether our original threshold of excluding scans which were rated as ‘Failed’ and including scans which were rated ‘Excellent’, ‘Good’ and ‘Doubtful’ was optimal, or whether additionally excluding scans rated as ‘Doubtful, resulted in a better prediction (a more stringent threshold resulting in many more exclusions according to manual ratings, N=207). We reran the analyses in step 2 by assessing how well the model predicted manual inclusion of rating ‘Excellent’ & ‘Good’ and exclusion for both rating ‘Doubtful’ & ‘Failed’. The 10-fold crossvalidation on the full dataset showed that AUC values were significantly lower using this more stringent threshold compared to the original threshold (mean AUC = .854; t(11.857) = −32.077, *p* = 6.9 × 10^−13^). Now, a total of 109 scans were misclassified (see Supplementary Figure 1), compared to 26 misclassifications with the original threshold. In addition, there were more scans classified with midrange (around 50) scan quality scores and the stability of the predictions was lower as can be seen from the higher variability of ratings between folds (see Supplementary Figure 1). This indicates that the original threshold was more optimal since the lower stability of the quality ratings with this more stringent threshold suggest that it is harder to distinguish between scans that should be excluded and included.

### Evaluation on novel datasets

In the last set of analyses we tested the external validity of our approach. First we used the full BrainTime dataset as the training set and the BESD dataset as the testing set (N = 112). When comparing the model-based binary classification of scan quality with manual ratings, the model had an accuracy of 0.893, sensitivity of 0.524 and specificity of 0.978. Ten scans were falsely included and two scans falsely excluded (total 10.7%). In order to reduce this error to 5%, we would recommend to manually check scans rated with a scan quality between 30% and 70%. The majority of misclassified scans will fall within these boundaries (see Figure 4 and Supplementary Figure 2).

We also used 10-fold cross validation on the BESD dataset to check whether rating of a subset of the data is sufficient to make reliable Qoala-T predictions. The mean AUC of the ten folds was AUC= 0.947 (SD = 0.011); results of all ten folds can be observed in Table 4. Post hoc t-tests showed that for the BESD compared to the BrainTime dataset mean AUC (*p* = 4.8 × 10^−6^), mean accuracy (*p* = 1.4 × 10^−4^), mean specificity (*p* = .001), and mean negative predictive value (*p* = 9.5 × 10^−5^) were significantly higher in the BrainTime dataset, whereas mean sensitivity (*p* = .37) and mean positive predictive value (*p* = .54) did not differ between these datasets. The mean number of falsely excluded scans was eleven (SD = 5.73) (~10%) and mean number of falsely included was three (SD = 1.60) (~3%). Although random forests are relatively robust to overfitting (Breiman, 2001; Strobl et al., 2009), this high SD and the statistical differences in predictive values compared to the subset-based model of the BrainTime data described earlier might indicate a tendency to overfit. Hence, the reliability of predictions probably increases with larger datasets or using larger subsets to rate manually. Another solution to prevent overfitting might be to further tune the hyperparameters in case of smaller datasets. For example, changing *mtry* to the amount of total variables increases the chance that the prediction model uses the most important predictor variables instead of randomly sampled suboptimal values (Bernard et al., 2009; Probst et al., 2018). Moreover, by setting a maximum to the amount of terminal nodes allowed (*maxnodes*) the model would not recursively try to fit data on trees with a very large depth (Chen & Ishwaran, 2012; Lin & Jeon, 2006).

**Table 4.** 10-fold cross validation using subsets of BESD data to predict manual quality ratings ofFreeSurfer-processed T1-weighted BrainTime scans

| Fold | AUC | Accuracy | Sensitivity | Specificity | PPV | NPV |
| --- | --- | --- | --- | --- | --- | --- |
| 1 | 0.944 | 0.810 | 0.895 | 0.790 | 0.500 | 0.970 |
| 2 | 0.961 | 0.822 | 0.895 | 0.805 | 0.515 | 0.971 |
| 3 | 0.957 | 0.930 | 0.778 | 0.963 | 0.824 | 0.952 |
| 4 | 0.941 | 0.802 | 0.947 | 0.768 | 0.486 | 0.984 |
| 5 | 0.923 | 0.802 | 0.842 | 0.793 | 0.485 | 0.956 |
| 6 | 0.943 | 0.871 | 0.842 | 0.878 | 0.615 | 0.960 |
| 7 | 0.957 | 0.901 | 0.842 | 0.915 | 0.696 | 0.962 |
| 8 | 0.955 | 0.881 | 0.789 | 0.902 | 0.652 | 0.949 |
| 9 | 0.944 | 0.911 | 0.789 | 0.939 | 0.750 | 0.951 |
| 10 | 0.949 | 0.911 | 0.947 | 0.902 | 0.692 | 0.987 |
| Mean | 0.947 | 0.864 | 0.857 | 0.866 | 0.622 | 0.964 |
| SD | 0.011 | 0.050 | 0.063 | 0.070 | 0.121 | 0.014 |

*Note.* AUC = area under the curve; PPV = positive predictive value (i.e., proportion of scans tested positive as exclusions that were correctly predicted to be excluded); NPV = negative predictive value (i.e., proportion of scans tested negative as exclusions that were correctly predicted to be included); SD = standard deviation.

We repeated the 10-fold cross validation on the BESD dataset by setting *mtry* to its maximum value (*mtry* = 185), which slightly increased the mean accuracy (0.880) and slightly reduced the standard deviation (SD = 0.040) compared to the values reported in Table 4. Additionally setting *maxnodes* to a value of 3 did not increase the mean accuracy (0.880) but slightly reduced the standard deviation somewhat more (SD = 0.019).

Finally, we used the full BrainTime dataset as the training set and data from the 13 ABIDE sites as the testing set (N = 760). When comparing the model-based binary classification of scan quality with manual ratings from the MRIQC team, the model had an accuracy of 0.809, a sensitivity of 0.783 and specificity of 0.815. Compared to the MRIQC manual ratings 30 scans were falsely included and 115 scans falsely excluded (total ~19%) by the Qoala-T model. The majority of scans misclassified by the Qoala-T model in the ABIDE dataset also fell within the 30%-70% boundaries of the Qoala-T score. Thus, most misclassified scans received Qoala-T scores that reflect higher uncertainty (i.e., around 50%), again suggesting that the amount of misclassifications can be reduced with an additional manual check of scans with scores between 30%-70%.

### Additional analyses: relation with age, sex, and MRIQC metrics

Linear regression revealed a significant relationship between Qoala-T scores and age in the BrainTime (*p* = 2 × 10^−16^), BESD (*p* = .016), and ABIDE (*p* = 4.4 × 10^−10^) datasets, showing that younger age is associated with lower scan quality (see Figure 7). Furthermore, in both the BrainTime (*p* = 4.3 × 10^−6^) and ABIDE (6.6 × 10^−4^) datasets, we found a significant relationship between Qoala-T scores and sex, showing that males had lower scan quality than females (the BESD sample is male only). Finally, linear regression analyses showed that Qoala-T scores were significantly related to MRIQC metrics in the BrainTime dataset (*F* (58,732) = 38.64, *p* = 2.2 × 10^−16^, *R^2^_ADJ_* = .736), BESD dataset (*F*(58,53) = 7.73, *p* = 1.8 × 10^−12^, *R^2^_ADJ_* = .779), and ABIDE dataset (*F*(64,694) = 18.68, *p* = 2.2 × 10^−16^, *R^2^_ADJ_* = .599).

**Figure 7.**
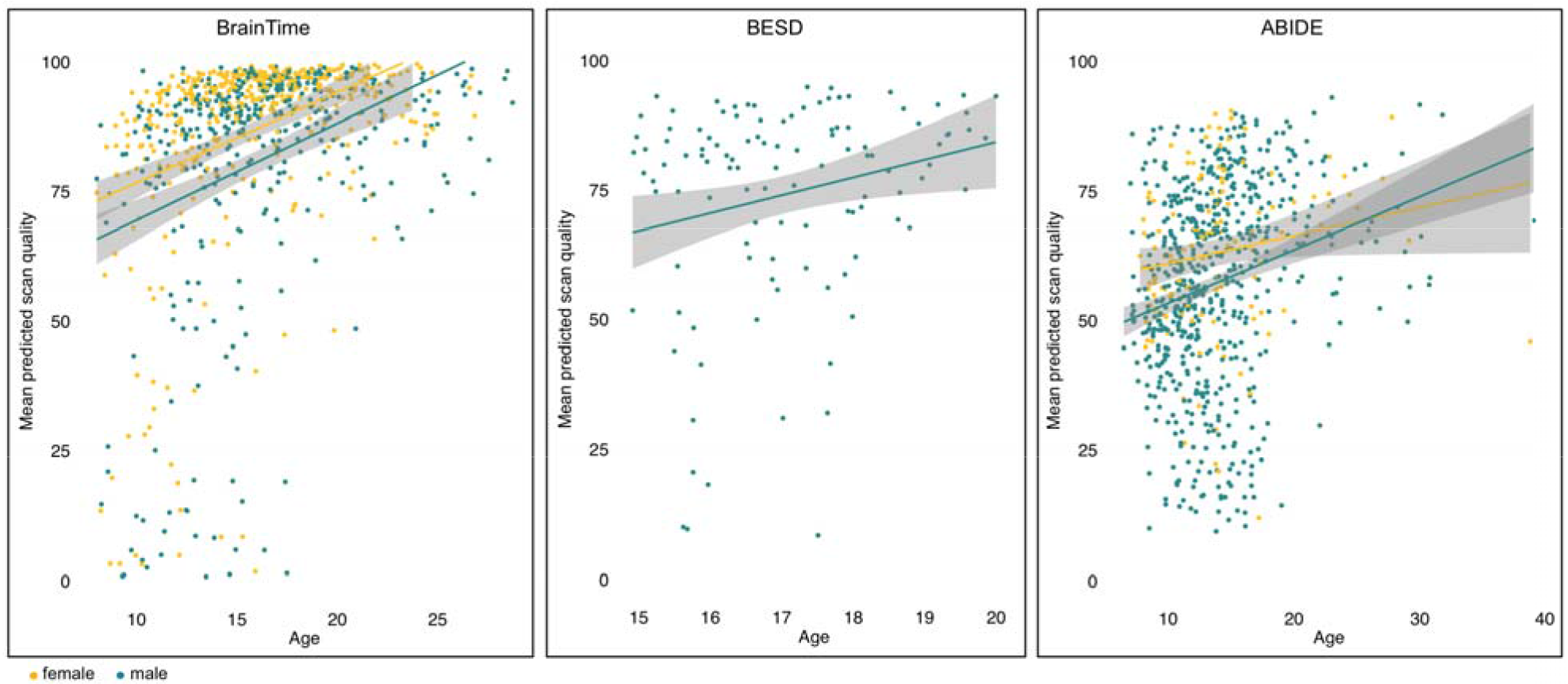
Predicted scan quality as a function of age and sex shown separately for the BrainTime, BESD and ABIDE datasets, showing younger participants and male participants were more likely to have lower scan quality. Age is displayed on the x-axis, mean values for predicted scan quality (0-100) are displayed on the y-axis, and colors indicate sex (greenish = male; yellow = female).

## Discussion

In the current study we present Qoala-T as a tool to automatically evaluate quality of FreeSurfer-processed structural MRI data. The modest agreement between different manual raters confirmed the subjectivity of manual QC ratings. We then showed that the random forests (RF) model was the best performing supervised-learning model. Manually assessed MRI quality could be predicted with both high sensitivity and specificity, by using data automatically segmented by FreeSurfer. In addition, we demonstrate that when only a relatively small amount of data is manually rated (in our case only 10% since we had a large dataset available), the Qoala-T tool is able to effectively predict the quality of the remaining data. Furthermore, the Qoala-T tool based on the BrainTime model can adequately predict the quality of unseen data. This automated procedure could therefore greatly help to reduce variability attached to manual QC (e.g., rater bias), thereby increasing the comparability of data quality between studies.

We looked for the best performing supervised-learning model that could distinguish FreeSurfer-processed scans that were manually rated with good quality from scans that were rated with insufficient quality. The model with the highest AUC values turned out to be the RF model, a supervised-learning technique suitable for analyzing high-dimensional data (Chen & Ishwaran, 2012; Strobl et al., 2009; Touw et al., 2013). This is in line with findings from a recent study that compared SVM and RF models for predicting MRI quality using raw data (Esteban et al., 2017). Our results show that using the RF model, we can predict manually assessed MRI quality with both high sensitivity (0.88) and specificity (0.97). This model also performed reasonably well on a smaller unseen dataset (BESD; sensitivity = 0.52; specificity = 0.98) and on a sample of comparable size from the ABIDE study (sensitivity = 0.78; specificity 0.82). Importantly, the Qoala-T model performed well even though the ABIDE manual ratings were online available ratings evaluated using a different procedure than used for the BrainTime model. This implies that the Qoala-T tool might be useful in addition to or in substitution for manual QC in other MRI studies using FreeSurfer. We therefore publicly share the Qoala-T source code through GitHub (https://github.com/Qoala-T/QC, allowing others to use the BrainTime model to generate Qoala-T scores in new datasets or to train the model using a subset of manually rated data. The use of such a validated and systematic QC procedure is of great importance for structural MRI studies, as the inclusion of data containing artifacts has shown to lead to spurious results, especially in developmental and clinical populations (Ducharme et al., 2016; Reuter et al., 2015).

Results of the current study also reflect the subjectivity of manual QC. We could establish only moderate inter-rater agreement, similar to a recent study that compared two raters (Esteban et al., 2017). This was the case even though in the current study QC was performed by five well-trained raters that followed the same detailed procedure. Hence, the results of the manual QC underline the need for objective and reliable QC methods by means of automated procedures. The automated Qoala-T tool that we propose may help overcome limitations of manual QC, including the subjectivity of these procedures and it may additionally reduce the time needed to perform QC. Note that although we aimed to reduce subjectivity bias, the current predictions are nevertheless based on manual QC. However, this bias is reduced (yet not fully removed) by determining the cutoff between inclusion and exclusion in a data driven way. This makes QC both less subjective and more comparable between studies. However, to achieve a full understanding of the quality of one,s dataset, experience in manual QC is still crucial.

In addition, this tool not considers raw image quality but the quality of subsequent segmentation, which is the data of interest in brain morphometric studies. A recent study found that image quality metrics (e.g., such as background voxels located outside the brain) derived from raw T1-weighted scans indeed affect FreeSurfer derived cortical thickness and surface area (White et al., 2018). However, raw image quality metrics typically use information from noise outside the head (Esteban et al., 2017; Pizarro et al., 2016; White et al., 2018). By using FreeSurfer derived measures from inside the brain we increase the likelihood of detecting artifacts that might remain unnoticed when they do not alter characteristics of the air around the head (Pizarro et al., 2016). Indeed, high quality segmentation is eventually conditional upon high quality of raw images. We demonstrate that the current FreeSurfer-based quality scores are consistent with raw image quality metrics from another QC tool (MRIQC), showing that the Qoala-T tool can be used both to check the quality of segmented MRI data and as a proxy for raw image quality and additional FreeSurfer derivatives (e.g., subcortical and cerebellar measures).

Manual QC often includes a multipoint scale as outcome variable, the present study for instance used a 4-point scale. It can be challenging to establish the right threshold for usable versus unusable scans. We therefore compared two different cutoffs for exclusion: one in which only scans rated as ‘Failed’ were excluded and one in which both scans rated as ‘Failed’ and ‘Doubtful’ were excluded. The results show that the predictive power of the model using the second, more stringent, threshold was significantly lower and led to much more misclassifications than the original threshold. As such the method applied in this paper can help identify the most optimal cutoff for scan exclusion.

We also found that the most important measures that predicted data quality in the random forests model were the left and right surface holes. These surfaces holes are FreeSurfer outputs that reflect topological defects in the surface reconstruction. Although the number of holes is related to the quality of the topological reconstruction, small-scale defects may still result in a usable reconstruction (Dale et al., 1999). Interestingly, a recent study found that the Euler number, which is directly related to the number of surface holes (Dale et al., 1999), was very useful in predicting data quality (Rosen et al., 2017). In that study, the Euler number was highly correlated with manual QC ratings, could discriminate unusable from usable scans with high accuracy, and could outperform other data quality measures such as those based on background voxels (Rosen et al., 2017). The predictions in the Qoala-T model are based on a combination of brain measures and not solely on surface hole measures, leading to increased model accuracy. Following the surface holes, the most important measure was the volume of white matter hypointensities. This measure might represent grey matter misclassified as white matter, and has been found to be higher in scans manually rated with bad compared to good and doubtful quality (Backhausen et al., 2016). Interestingly most of the other variables with high importancy values were measures of cortical thickness. This could be due to a greater sensitivity of thickness estimates to motion artifacts compared to surface area, as a previous study found associations between motion and thickness in the absence of associations between motion and surface area (Alexander-Bloch et al., 2016).

Exploratory analyses of age and sex effects confirmed previous findings showing that younger participants and male participants are more likely to have lower scan quality (Ducharme et al., 2016; Pardoe et al., 2016; White et al., 2018). These findings once again underline the importance of taking scan quality into account in developmental studies, as scan quality might confound findings attributed to age effects. One possible solution could be to use measures of scan quality such as the Qoala-T score as a covariate when studying age effects, in addition to using such measures as guidance for participant exclusion (see also recommendations by White et al., 2018).

### Practical Recommendations

The Qoala-T source code including instructions is publicly available through GitHub (https://github.com/Qoala-T/QC) and we provide an R Shiny app through which the Qoala-T model can be run with a few mouse clicks without having R installed (https://qoala-t.shinyapps.io/qoala-t_app/). Users can choose to either predict scan quality by using the BrainTime model (which does not necessarily require manual QC) or to predict scan quality by building a model based on manual ratings on a subset of their own data. Both options result in a Qoala-T score (ranging from 0 to 100) for every individual scan. This score is based on class probability, with higher numbers indicating a higher chance of being a high-quality scan, and thus a higher likelihood for the scan to be included. To reduce misclassification we would recommend to manually check scans rated with a scan quality between 30% and 70%, since many misclassified scans will fall within these boundaries (see Figure 4 and Supplementary Figure 2). If users prefer to use the Qoala-T tool on a manually rated subset of their own data to predict the quality of the remaining data, we would recommend using a sufficient number of manually rated scans (e.g., 10 % in larger datasets, but at least 50 scans in smaller datasets, of which around >10% were rated Failed) to be sure the model is fed with sufficient variability in scan quality (see for example the difference in variability in output variables between the folds in Tables 3 and 4). Furthermore, when using smaller datasets to predict scan quality by rating an own-data subset (i.e., not using the BrainTime model) we recommend considering additional tuning of the hyperparameters (e.g., *mtry* and *maxnodes*, see Evaluation on novel datasets in the Results section), as this might improve performance and could reduce overfitting. In general, smaller datasets are less suited to predict scan quality by rating a subset that also can be seen from the significantly lower predictive values when using 10% of the data within the BESD sample (N = 112) compared to using 10% of the data within the BrainTime sample (N = 784).

In addition to increasing comparability between studies, our tool could potentially save time by performing manual QC on only a part of the dataset. Using our protocol, manual QC took about 5-10 minutes per scan. By examining only scans rated with a scan quality between 30% and 70% this would lead to more than 60.8 hours reduced out of 65.3 total hours for the BrainTime dataset (729 out of 784 scans fell outside the 30%-70% range), 7.2 hours reduced out of 9.3 hours for the BESD dataset (86 out of 112 scans fell outside the 30%-70% range), and 24.1 hours reduced out of 63.3 hours for the ABIDE dataset (289 out of 760 scans fell outside the 30%-70% range). Given the current state of the art, it is not advised to fully replace manual efforts by automatic tools since Qoala-T and other automated QC measures (e.g., Esteban et al., 2017; Rosen et al., 2017; White et al., 2018) currently show no perfect overlap with manual QC ratings (note however that this may be due to errors in either automated or manual classification). Finally, the Qoala-T tool could be complemented by tools such as Mindcontrol (Keshavan et al., 2018a) that could be used to edit scans with fair quality scores but with errors in segmentation.

### Limitations

Several limitations of the current study deserve consideration. First, in the current study we used data from one scanner site to train the model. Previous studies have shown that the accuracy of data quality predictions can vary across sites (Esteban et al., 2017; Rosen et al., 2017; White et al., 2018). We found that accuracy was indeed lower but still acceptable when predicting quality of scans from different ABIDE sites compared to those acquired at the same site. Second, the Qoala-T tool requires FreeSurfer-processed scans, which limits use of this procedure to one software package that also requires extensive processing. However, FreeSurfer is a widely used platform; researchers that are already planning to use FreeSurfer can easily use the Qoala-T tool with only a few additional steps. Third, since the manual QC rating is the outcome measure of our model, these manual scores will remain the principle QC rating. As such, we cannot distinguish the nature of misclassifications. These could be related to human error or classification error of the model. We should note, however, that the current inter-rater reliability of manual raters is relatively low. Although this might accurately reflect reality in which many scans fall in the subjective gray area of QC, this also means that the standard on which the Qoala-T model is trained could be improved. However, the goal of the current study was not to establish a new gold standard based on manual QC, but to reduce bias in QC by jointly using automated and manual QC. In this way, data driven quality scores can inform the final decisions about quality and this will help to make QC process more objective. Future studies could also combine the current method with crowdsourcing methods to account for individual sensitivity and specificity of multiple manual raters (Raykar et al., 2010; Warfield et al., 2004). See also Keshavan et al. (2018b) for a recent example of using ratings from over 250 citizen scientists for brain imaging QC. Fourth, the results of the Qoala-T model show both good sensitivity and specificity to detect which scans are of sufficient quality, with a somewhat higher specificity. Ideally however, the prediction model should have near perfect sensitivity or specificity, such that no manual recheck is necessary for either the scans predicted to be included or to be excluded. With the current model we advise users instead to focus manual QC on the scans with midrange probabilities, where the model is less certain about inclusion or exclusion. Fifth, although the main aim of the Qoala-T tool is to increase comparability of QC between studies, we maintained options to facilitate flexible usage that come at the cost of full standardization (e.g., others can train the model on a subset of their data and different rating procedures can be adapted to study needs). Even when the model will be trained on different data or rating procedures, comparability is already considerably increased compared to the current practice in which many studies at best report very generally about QC procedures (Backhausen et al., 2016).

### Conclusions

In sum, we demonstrate that the Qoala-T tool is a useful tool to assess quality of MRI data that is automatically segmented using FreeSurfer. When a subset of data has been manually rated, the Qoala-T tool is able to predict the quality of the remaining data with high accuracy. Moreover, the quality of novel datasets can also be adequately predicted using the Qoala-T tool. We have made this tool publicly available, such that other researchers can use it as an add-on to manual QC in their studies. This could greatly increase comparability of data quality between studies.

## Supporting information

Supplementary Materials v3 final

## Acknowledgements

The BrainTime study was funded by a starting grant of the European Research Council (ERC-2010-StG-263234 awarded to Eveline A. Crone). We are grateful to all participants and their parents for their willingness to participate. For the use of the BESD dataset, the authors like to thank all participants and their parents, the participating centers (De Jutters Palmhuis Forensic Psychiatric Unit, Forensic Center and Correctional Facility Teylingereind, Centrum Autisme Rivierduinen, Curium-LUMC), Moji Aghajani, Romy Emmerig and Simone van Montfort for their help with data collection and to all involved in the design of this study (Moji Aghajani, Olivier Colins, Natasja van Lang, Arne Popma, Robert Vermeiren, Nic van der Wee). The BESD study was supported by the Netherlands Organization for Scientific Research (NWO) Grant No. 056-23-011. We also thank the individual ABIDE sites that collected data and all participants who participated in their studies. We like to thank all ABIDE investigators for their effort to make these data publicly available. Finally, we thank Michel Villerius for help with running Singularity containers on the LUMC Shark computer cluster.

