## Supplementary Materials v3 final for "Qoala-T: A supervised-learning tool for quality control of FreeSurfer segmented MRI data"

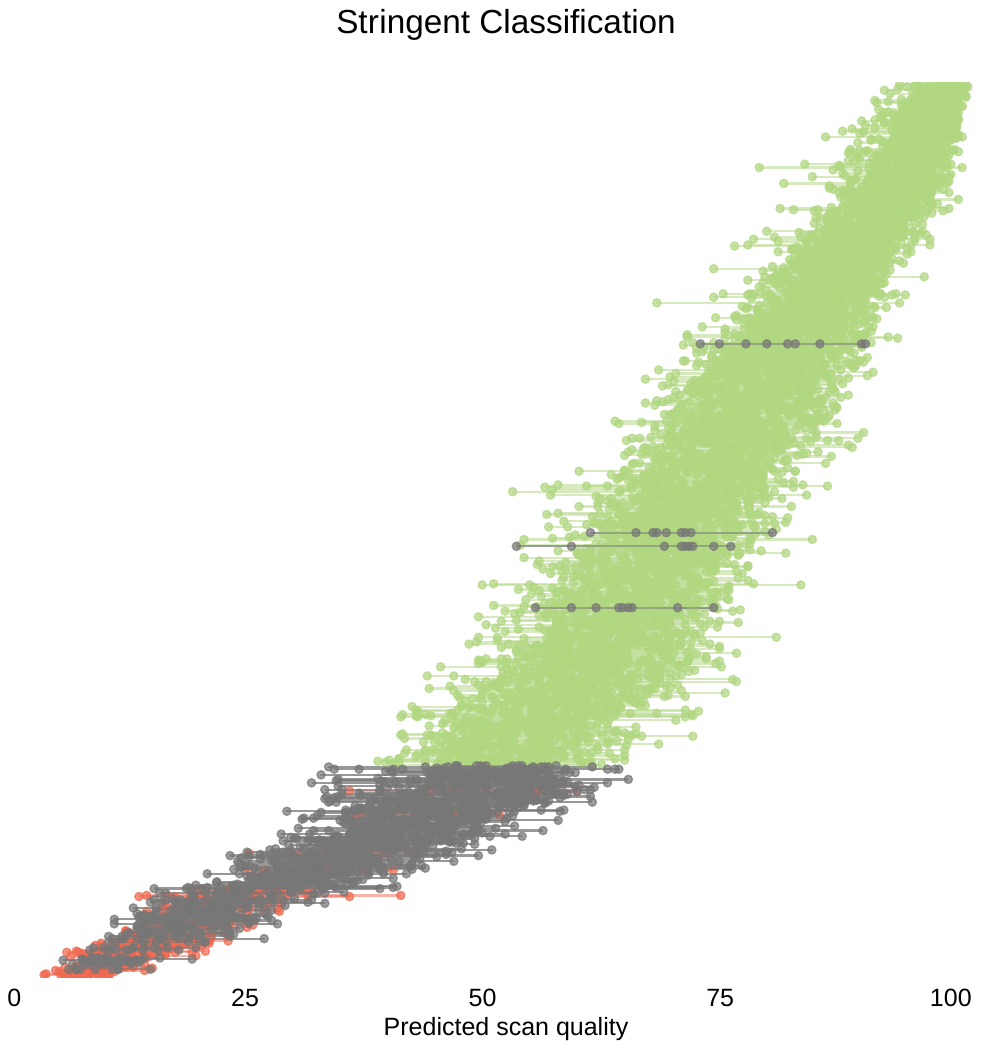

*Supplementary Figure 1. Stringent* *classification (both ‘Doubtful’ and ‘Failed’ scans excluded), showing larger prediction errors and more uncertainty ratings for estimated scan quality (x-axis). Scans with an average scan quality <50% are predicted to be exclude (right), and vice versa scans with an average scan quality > 50% are predicted to be included (left). Colors indicate goodness of predicted classification (green = 'correct include', red = 'correct exclude', grey = 'incorrect classification'). A total of 149 scans were misclassified.*

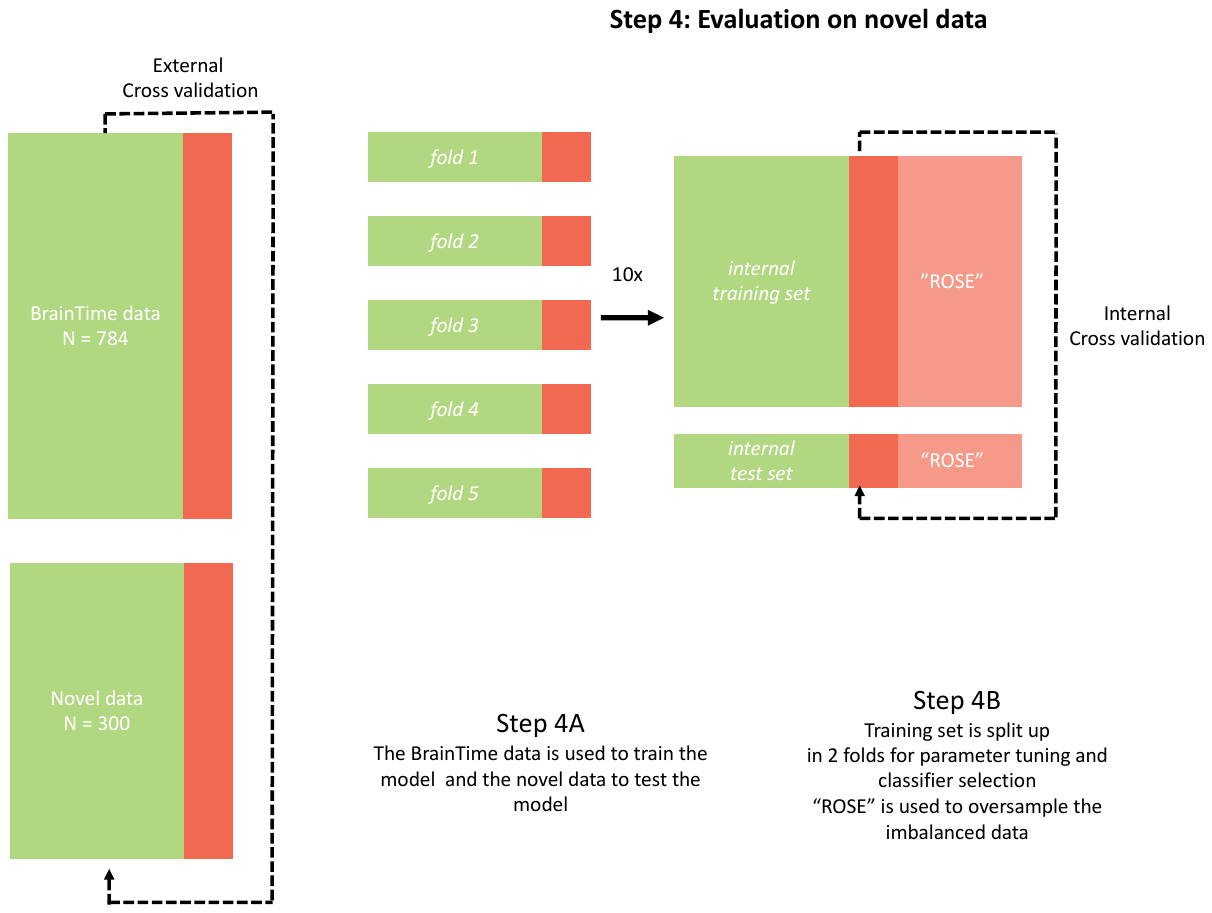

*Supplementary Figure 2. Illustration of Step 4 (Evaluation on novel data) for the development of the Qoala-T tool. Green indicates included scans, red indicates scans to be excluded.*

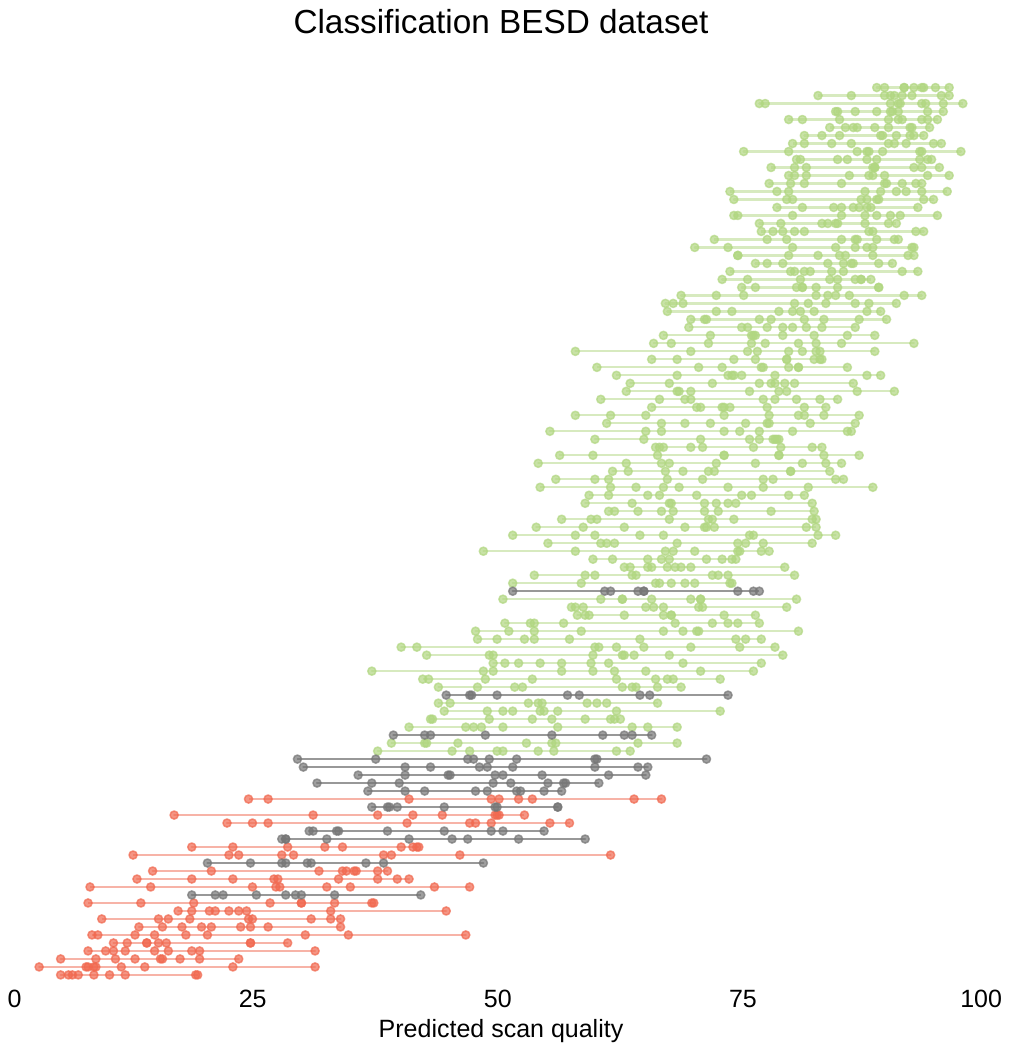

*Supplementary Figure 3. Predicted scan quality for the BESD dataset. For each scan 9 values for predicted scan quality (0-100) are displayed on the x-axis. Each value on the y-axis represents a scan ordered by mean scan quality. Lines between dots connect 9 values of predicted scan quality for a single scan. Scans with an average scan quality <50% are predicted to be excluded, and scan with an average scan quality >50% are predicted to be included. Colors indicate goodness of predicted classification (green = 'correct include', red = 'correct exclude', grey = 'incorrect classification'). A total of 13 scans out of a total of 112 scans (11.6%) were misclassified.*

*Supplementary Table 1. Mean permutation importance of explanatory variables for the prediction of scan quality for the 10 folds (see also Figure 6 for a graphical representation of the top 50 highest importance values). The importance measure represents the mean increase in classification error that results from randomly permuting that variable. Note that the importance value is a relative value that allows for a quick assessment of the relevance of a predictor for the outcome of interest.*

| **Structure name** | **Mean** | **SD** |
| --- | --- | --- |
| rhSurfaceHoles | 96.89 | 3.23 |
| lhSurfaceHoles | 94.85 | 8.92 |
| WM.hypointensities | 57.07 | 6.40 |
| rh_entorhinal_thickness | 41.51 | 9.87 |
| lh_precentral_thickness | 39.79 | 9.59 |
| rh_precentral_thickness | 31.27 | 7.22 |
| rh_caudalmiddlefrontal_thickness | 30.83 | 6.05 |
| lh_pericalcarine_thickness | 27.05 | 6.03 |
| rh_pericalcarine_thickness | 26.07 | 4.64 |
| lh_caudalmiddlefrontal_thickness | 23.81 | 5.44 |
| rh_superiortemporal_thickness | 23.29 | 6.81 |
| lh_entorhinal_thickness | 22.99 | 4.79 |
| rh_supramarginal_thickness | 21.39 | 5.39 |
| lh_superiorfrontal_thickness | 16.93 | 3.25 |
| lh_superiortemporal_thickness | 15.52 | 5.26 |
| lh_lingual_thickness | 14.68 | 4.45 |
| X5th.Ventricle | 14.12 | 2.57 |
| rh_insula_thickness | 13.13 | 4.27 |
| lh_superiorparietal_area | 12.02 | 4.47 |
| rh_precentral_area | 10.93 | 4.86 |
| lh_supramarginal_thickness | 9.91 | 4.67 |
| lh_lateralorbitofrontal_thickness | 9.05 | 3.23 |
| lh_precentral_area | 8.11 | 3.76 |
| rh_middletemporal_thickness | 8.05 | 3.45 |
| lh_rostralmiddlefrontal_thickness | 7.96 | 2.77 |
| lh_postcentral_area | 7.66 | 2.50 |
| lh_middletemporal_thickness | 7.65 | 2.94 |
| rh_lingual_thickness | 7.63 | 2.73 |
| lh_insula_thickness | 7.59 | 3.21 |
| lh_inferiortemporal_thickness | 7.39 | 2.95 |
| lh_parsopercularis_thickness | 7.14 | 2.27 |
| rh_supramarginal_area | 7.07 | 3.18 |
| lh_superiorfrontal_area | 6.96 | 4.57 |
| rh_rostralmiddlefrontal_thickness | 6.83 | 2.85 |
| rh_superiorfrontal_thickness | 6.82 | 2.00 |
| rh_lateralorbitofrontal_thickness | 6.61 | 4.91 |
| rh_inferiorparietal_area | 6.12 | 2.64 |
| rh_inferiorparietal_thickness | 5.95 | 2.96 |
| Right.VentralDC | 5.92 | 1.14 |
| rh_rostralanteriorcingulate_area | 5.56 | 1.95 |
| rh_inferiortemporal_thickness | 5.50 | 2.06 |
| rh_superiorfrontal_area | 5.08 | 1.41 |
| rh_postcentral_area | 5.07 | 1.68 |
| Left.Thalamus.Proper | 5.01 | 1.73 |
| lh_parsorbitalis_thickness | 4.85 | 2.25 |
| Right.Cerebellum.White.Matter | 4.79 | 1.46 |
| Left.VentralDC | 4.78 | 1.49 |
| rh_MeanThickness_thickness | 4.72 | 1.38 |
| rh_superiorparietal_area | 4.41 | 1.89 |
| lh_cuneus_thickness | 4.40 | 2.17 |
| rh_entorhinal_area | 4.30 | 1.63 |
| rh_parsopercularis_thickness | 4.00 | 1.59 |
| lh_MeanThickness_thickness | 3.98 | 1.80 |
| rh_caudalmiddlefrontal_area | 3.86 | 1.43 |
| lh_inferiorparietal_thickness | 3.79 | 1.93 |
| rh_paracentral_thickness | 3.75 | 1.73 |
| Brain.Stem | 3.73 | 0.82 |
| lh_supramarginal_area | 3.67 | 1.78 |
| lh_insula_area | 3.63 | 1.47 |
| lh_isthmuscingulate_thickness | 3.63 | 1.58 |
| lh_rostralmiddlefrontal_area | 3.63 | 2.37 |
| rh_medialorbitofrontal_thickness | 3.60 | 1.74 |
| lh_superiortemporal_area | 3.42 | 1.31 |
| lh_inferiorparietal_area | 3.42 | 1.98 |
| rh_rostralmiddlefrontal_area | 3.40 | 0.97 |
| rh_cuneus_thickness | 3.38 | 1.14 |
| lh_precuneus_thickness | 3.31 | 1.24 |
| CSF | 3.23 | 1.36 |
| rh_paracentral_area | 3.17 | 2.11 |
| lh_transversetemporal_area | 3.13 | 1.41 |
| rh_transversetemporal_area | 3.09 | 1.48 |
| rh_lateralorbitofrontal_area | 3.06 | 0.99 |
| lh_parstriangularis_thickness | 3.03 | 1.07 |
| rh_inferiortemporal_area | 2.96 | 0.72 |
| lh_medialorbitofrontal_thickness | 2.84 | 0.94 |
| lh_caudalmiddlefrontal_area | 2.81 | 1.53 |
| rh_WhiteSurfArea_area | 2.79 | 1.37 |
| rh_rostralanteriorcingulate_thickness | 2.74 | 0.97 |
| lh_postcentral_thickness | 2.72 | 1.81 |
| BrainSegVol.to.eTIV | 2.71 | 0.92 |
| lh_WhiteSurfArea_area | 2.70 | 1.58 |
| X3rd.Ventricle | 2.67 | 0.74 |
| Right.Pallidum | 2.65 | 1.23 |
| rh_lingual_area | 2.59 | 1.41 |
| lh_parstriangularis_area | 2.59 | 1.10 |
| lh_posteriorcingulate_thickness | 2.52 | 1.05 |
| Left.Caudate | 2.48 | 1.47 |
| rh_superiortemporal_area | 2.47 | 1.51 |
| rh_precuneus_thickness | 2.45 | 0.82 |
| lh_paracentral_area | 2.41 | 1.09 |
| rh_medialorbitofrontal_area | 2.31 | 1.35 |
| Optic.Chiasm | 2.31 | 0.97 |
| rh_middletemporal_area | 2.23 | 1.14 |
| Left.vessel | 2.09 | 0.94 |
| CortexVol | 2.09 | 1.06 |
| lh_medialorbitofrontal_area | 2.08 | 1.12 |
| Left.Cerebellum.White.Matter | 2.01 | 0.65 |
| SupraTentorialVolNotVent | 1.99 | 0.79 |
| CC_Mid_Anterior | 1.99 | 0.60 |
| lh_rostralanteriorcingulate_thickness | 1.97 | 0.89 |
| X4th.Ventricle | 1.93 | 0.87 |
| lhCortexVol | 1.89 | 1.02 |
| rh_parsorbitalis_thickness | 1.88 | 0.97 |
| rhCortexVol | 1.82 | 0.85 |
| Left.Accumbens.area | 1.77 | 1.02 |
| lh_paracentral_thickness | 1.74 | 0.86 |
| rh_pericalcarine_area | 1.74 | 1.04 |
| BrainSegVolNotVentSurf | 1.73 | 0.49 |
| SubCortGrayVol | 1.73 | 0.66 |
| Right.Accumbens.area | 1.72 | 0.87 |
| lh_lateralorbitofrontal_area | 1.72 | 0.96 |
| rh_parstriangularis_thickness | 1.67 | 0.41 |
| EstimatedTotalIntraCranialVol | 1.66 | 0.66 |
| Right.vessel | 1.65 | 1.13 |
| BrainSegVolNotVent | 1.63 | 0.77 |
| Right.Caudate | 1.62 | 1.08 |
| lh_parsopercularis_area | 1.62 | 0.71 |
| TotalGrayVol | 1.62 | 0.66 |
| Right.Thalamus.Proper | 1.59 | 0.80 |
| SupraTentorialVolNotVentVox | 1.55 | 0.50 |
| SupraTentorialVol | 1.53 | 0.55 |
| Right.Lateral.Ventricle | 1.49 | 0.65 |
| CC_Posterior | 1.47 | 0.58 |
| lh_lateraloccipital_area | 1.46 | 0.48 |
| BrainSegVol | 1.41 | 0.94 |
| Right.choroid.plexus | 1.41 | 0.53 |
| lh_fusiform_thickness | 1.41 | 0.60 |
| lh_lingual_area | 1.40 | 1.18 |
| Left.choroid.plexus | 1.39 | 0.78 |
| lh_posteriorcingulate_area | 1.37 | 1.47 |
| rh_posteriorcingulate_thickness | 1.37 | 0.49 |
| lh_lateraloccipital_thickness | 1.34 | 0.87 |
| rh_postcentral_thickness | 1.33 | 0.56 |
| lh_superiorparietal_thickness | 1.32 | 0.46 |
| MaskVol | 1.32 | 0.54 |
| CC_Anterior | 1.30 | 0.56 |
| lh_entorhinal_area | 1.28 | 0.41 |
| rh_parsopercularis_area | 1.26 | 0.76 |
| lh_rostralanteriorcingulate_area | 1.23 | 0.98 |
| rh_transversetemporal_thickness | 1.23 | 0.55 |
| lh_precuneus_area | 1.22 | 0.69 |
| Left.Lateral.Ventricle | 1.22 | 0.63 |
| rh_parahippocampal_thickness | 1.19 | 0.44 |
| lh_transversetemporal_thickness | 1.16 | 0.53 |
| Right.Inf.Lat.Vent | 1.14 | 0.54 |
| lh_isthmuscingulate_area | 1.13 | 0.41 |
| Right.Cerebellum.Cortex | 1.12 | 0.51 |
| rh_superiorparietal_thickness | 1.12 | 0.44 |
| rh_isthmuscingulate_thickness | 1.12 | 0.66 |
| lh_parahippocampal_area | 1.11 | 0.39 |
| rh_precuneus_area | 1.11 | 0.45 |
| lh_parahippocampal_thickness | 1.09 | 0.33 |
| rh_caudalanteriorcingulate_area | 1.09 | 0.30 |
| Left.Putamen | 1.08 | 0.58 |
| rh_parahippocampal_area | 1.06 | 0.66 |
| lh_parsorbitalis_area | 1.05 | 0.34 |
| rh_fusiform_thickness | 1.05 | 0.55 |
| Left.Amygdala | 1.05 | 0.38 |
| lh_inferiortemporal_area | 1.04 | 0.39 |
| CC_Mid_Posterior | 0.93 | 0.30 |
| lh_caudalanteriorcingulate_thickness | 0.93 | 0.59 |
| rh_caudalanteriorcingulate_thickness | 0.92 | 0.43 |
| Right.Amygdala | 0.91 | 0.29 |
| CC_Central | 0.91 | 0.33 |
| Left.Cerebellum.Cortex | 0.90 | 0.64 |
| rh_lateraloccipital_area | 0.89 | 0.37 |
| Left.Hippocampus | 0.88 | 0.23 |
| Right.Hippocampus | 0.87 | 0.47 |
| rh_parstriangularis_area | 0.87 | 0.39 |
| rh_parsorbitalis_area | 0.87 | 0.48 |
| lh_pericalcarine_area | 0.87 | 0.43 |
| rh_insula_area | 0.86 | 0.73 |
| rh_lateraloccipital_thickness | 0.85 | 0.56 |
| Left.Inf.Lat.Vent | 0.85 | 0.60 |
| Right.Putamen | 0.78 | 0.34 |
| rh_fusiform_area | 0.69 | 0.39 |
| Left.Pallidum | 0.69 | 0.38 |
| rh_isthmuscingulate_area | 0.68 | 0.41 |
| rh_cuneus_area | 0.66 | 0.26 |
| MaskVol.to.eTIV | 0.65 | 0.49 |
| rh_posteriorcingulate_area | 0.63 | 0.29 |
| lh_caudalanteriorcingulate_area | 0.62 | 0.31 |
| lh_middletemporal_area | 0.61 | 0.27 |
| lh_fusiform_area | 0.47 | 0.44 |
| lh_cuneus_area | 0.41 | 0.37 |

*Supplementary Table 2. Distribution of manual QC labels for the different datasets. Note that for the ABIDE dataset manual ratings released by the MRIQC project were used (https://github.com/poldracklab/mriqc; Esteban et al., 2017). These scans were rated on a three-point scale (accept/doubtful/exclude).*

|  | BrainTime | BESD | *ABIDE* |
| --- | --- | --- | --- |
| Excellent | 214 | 19 | *608 (accept)* |
| Good | 363 | 51 |  |
| Doubtful | 161 | 21 | *14* |
| Failed | 46 | 21 | *138 (exclude)* |
| Total | 784 | 112 | *760* |

*Supplementary Table 3. List of image quality metrics used to assess relations between Qoala-T scores and MRIQC metrics. For more information on these metrics see Esteban et al. (2017) and* [*https://mriqc.readthedocs.io/en/stable/iqms/t1w.html*](https://mriqc.readthedocs.io/en/stable/iqms/t1w.html)*. Asterisks indicate significant relation with Qoala-T scores in the different datasets: *^BT^ = BrainTime, *^BESD^ = BESD, *^ABIDE^ = ABIDE.*

| Abbreviation | Metric |
| --- | --- |
| CJV | Coefficient of joint variation*^BT^ |
| CNR | Contrast-to-noise ratio |
| EFC | Entropy focus criterion*^ABIDE^ |
| FBER | Foreground-to-background energy ratio |
| FWHM | Full width at half maximum (average*^BT^, x*^BT^, y, and z) |
| ICV | Intracranial volume fractions of CSF*^BESD^, GM*^BT, BESD^ and WM |
| INU | Intensity non-uniformity (range*^ABIDE^ and median*^BT^) |
| QI1 | Mortamet’s quality index 1 |
| QI2 | Mortamet’s quality index 2 |
| RPVE | Residual partial volume effects of CSF*^BT^, GM*^BESD^ and WM*^BT, BESD^ |
| SNR | Signal-to-noise ratio of CSF*^BT, BESD, ABIDE^, GM*^BT^, WM, and total*^BT^ |
| SNRd | Dietrich’s SNR of CSF*^BESD^, ^ABIDE^, GM*^BT, BESD^, WM, and total*^BT, BESD, ABIDE^ |
| SSTATS | Mean, standard deviation, 5% percentile and 95% percentile of the distribution of background*^BT, BESD, ABIDE^, CSF*^BT, ABIDE^, GM*^BT, BESD^ and WM*^BT, BESD, ABIDE^ |
| TPM | Overlap of tissue probability maps of CSF, GM*^BT, ABIDE^ and WM*^ABIDE^ |
| WM2MAX | White-matter to maximum intensity ratio*^BT^ |

**Manual quality control procedure for structural T1-weighted scans processed in FreeSurfer**

Brain and Development Research Center, Leiden University

October 2017, revised July 2018

### Background information

Manual quality check (QC) procedures of T1-weigthed MRI scans are susceptible to both inter-rater and intra-rater variability, and the methods used differ across studies and sites. With this manual we aim to provide more objective guidelines for the quality check of T1-weighted scans. Therefore, a step-by-step approach of our in house developed manual QC procedure is provided in this protocol. In addition, examples of MRI scans that we consider of sufficient and insufficient quality are included.

Tissue classification and anatomical labeling was performed on the basis of the T1- weighted scan using FreeSurfer v6.0.0 software (http://surfer.nmr.mgh.harvard.edu/). For the quality assessment described in this manual the gui **TkMedit** was used to visually inspect all FreeSurfer-processed scans (the end product of the pre-processing FreeSurfer pipeline). For more background information and tutorials on these processing steps in FreeSurfer, visit the website <https://surfer.nmr.mgh.harvard.edu/>.

In this manual, we discuss our manual quality control procedure of FreeSurfer-processed T1-weighted anatomical scans. After data collection, we processed the data as described in the Method section of the manuscript (see preprint at <https://doi.org/10.1101/278358>), resulting in FreeSurfer-processed scans for each participant. These FreeSurfer-processed scans were manually controlled for quality.

### 1. Set up the environment for quality control procedure

Prior to this step, the scans for each participant have been pre-processed in FreeSurfer (<http://freesurfer.net/fswiki/FreeSurferAnalysisPipelineOverview>).

**1.1 Start FreeSurfer and tkmedit (in Linux environment)**

To start quality control procedure for a specific scan from a participant, open FreeSurfer:

module load freesurfer/6.0.0

Set up path for your subject directory:

export SUBJECTS_DIR=/[subjects folder]/

After this set-up, the following steps can be repeated for each individual scan. The code below opens the brainmask, the original T1-weighted scan as auxiliary image and overlays the reconstructed pial and gm/wm boarder surfaces of a processed T1-weigthed scan:

tkmedit $CorrectDir brainmask.mgz -aux T1.mgz -surface lh.white -aux-surface rh.white

Below is an example of a script that automatizes this process, only the scan ID has to be entered.

echo "Please enter scan ID”

read scanID

echo "Starting quality control for {scanID}"

for dir in /$SUBJECTS_DIR /*/;

do

if [[ $dir == *"{scanID}"* ]]

then

CorrectDir=${dir#/$SUBJECTS_DIR/}

fi

done

view="tkmedit $CorrectDir brainmask.mgz -surfs -aux wm.mgz

eval $view

This code opens an image of the parcellated brain in TkMedit (see Figure M1) (<https://surfer.nmr.mgh.harvard.edu/fswiki/TkMeditGuide>). Two surfaces are displayed: the pial surface is indicated in red and the division between grey/white matter (GM/WM) boarder is indicated in yellow. The original T1-weigthed scan (including skull) is opened as auxiliary file; you can easily switch to this scan using the TkMedit gui (button 4, see Figure M2).

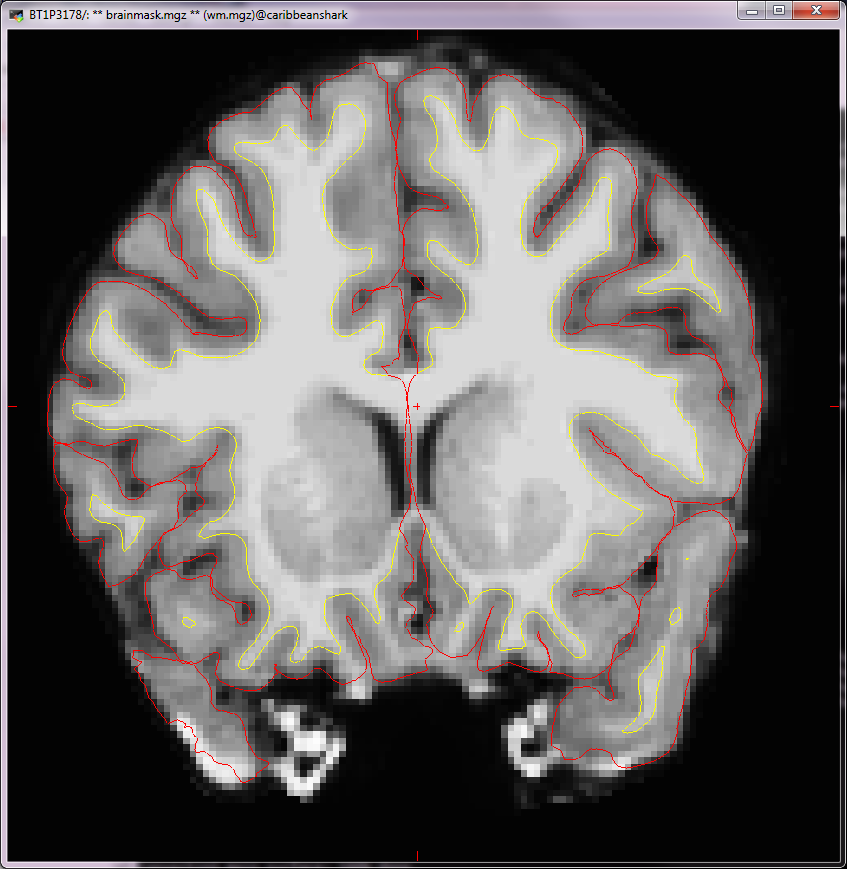

**Figure M1.** Example of parcellated scan opened in tkmedit, with pial surface (red) and WM/GM division (yellow).

**1.2 Set-up TkMedit window**

For quality control, we scroll through the anterior to the posterior regions of the brain. The scan will automatically open at slice 128, so to start your quality control procedure in the anterior region of the brain: either type “215” in box 1 (see Figure M2), scroll to the anterior regions using the “+” button, or type “Slice 215” in the command line. Next, use the “-“ button to scroll towards the posterior part of the brain in a slow pace, or use the arrow keys. To ensure comparability between subjects, make sure that when you enlarge the TkMedit window it is always approximately the same size for every subject. You can also zoom in on the brain (if necessary) (see button in TkMedit interface shown in Figure M2.2). In addition, you can activate the cursor so you can move the image around to inspect relevant parts (button in TkMedit interface shown in Figure M2.3).

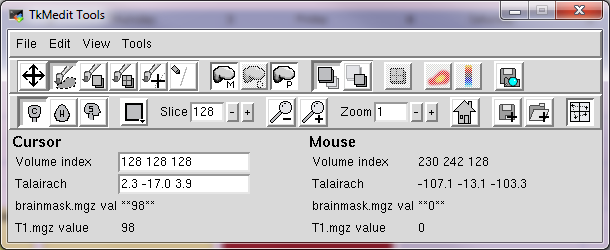

**4**

**2**

**3**

**1**

Figure M2. Example of TkMedit interface, with highlighted areas indicating slice selector (1), zoom in button (2), cursor activation button (3) and auxiliary file activation button (4).

### 2. Quality control procedure

**2.1 General information quality control procedure**

After setting up the environment for the quality control procedure, each T1-weigthed scan has to be screened carefully. Four criteria are used to assess scan quality:

1) Is the reconstructed image affected by movement?
2) Is (part of) the temporal pole missing in the reconstruction?
3) Is non-brain tissue (e.g. dura/skull) included in the reconstruction of pial surface (red line)?
4) Are parts of the cortex missing in the reconstruction (other than temporal poles)?

For each criterion 0 is entered when the answer is ‘No, no errors are visible’, 1 is entered when the answer is ‘Yes, errors are visible’. We only score 1 if the criterion is met in at least 3 consecutive slices.

Each criterion has to be assessed for the left and right hemisphere separately. As such, we inspect each MRI scan twice (e.g. first scrolling back and forth through the left hemisphere, then repeating the procedure for the right hemisphere). Together, these criteria (more details are provided below) lead to a final evaluation of the scan, expressed in a numerical rating of overall quality (1-4: Excellent (1), Good (2), Doubtful (3), Failed (4)).

**2.2 Keeping track of QC**

To keep track of QC for each T1-weigthed scan, we used a datafile (see template on https://github.com/Qoala-T/QC/blob/master/Template_ManualQC_Qoala-T.xlsx). Each line contains information on a single T1-weigthed scan, where each quality criteria is represented in the columns (see Figure M3):

Column A (*Subject*): the unique subject/scan number

Column B (*Final Score*) the final rating for an individual scan

Columns C-M: 4 criteria for both left and right hemispheres (LH = left hemisphere, RH = right hemisphere)

Column N (*Notes)*: additional comments

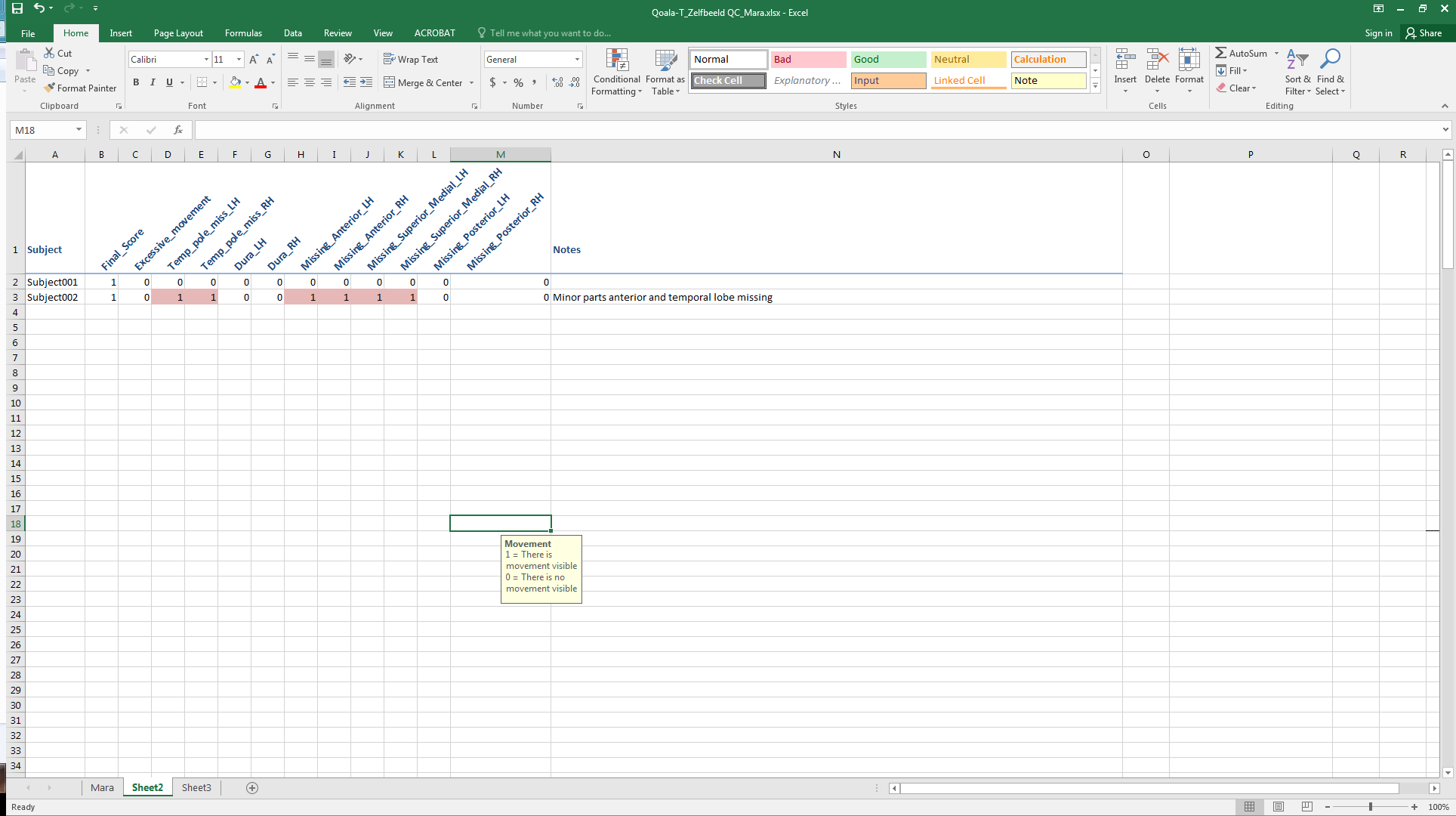

Figure M3. Example of quality control file, with individual lines with information for each scan.

.**2.3 Criteria for quality control**: **step-by-step**

Below we provide step-by-step details of the four quality criteria.

Criterion 1: Is the reconstructed image affected by movement?
*(0=NO, 1=YES)*

If participants have moved too much in the scanner (Backhausen et al., 2016), this is sometimes visible as rings (like growth rings on a tree; see Figure M4 for an example). Even though reconstruction has been completed, it is clearly not a good reconstruction. *Excessive movement* (column C) is therefore scored with “1”.

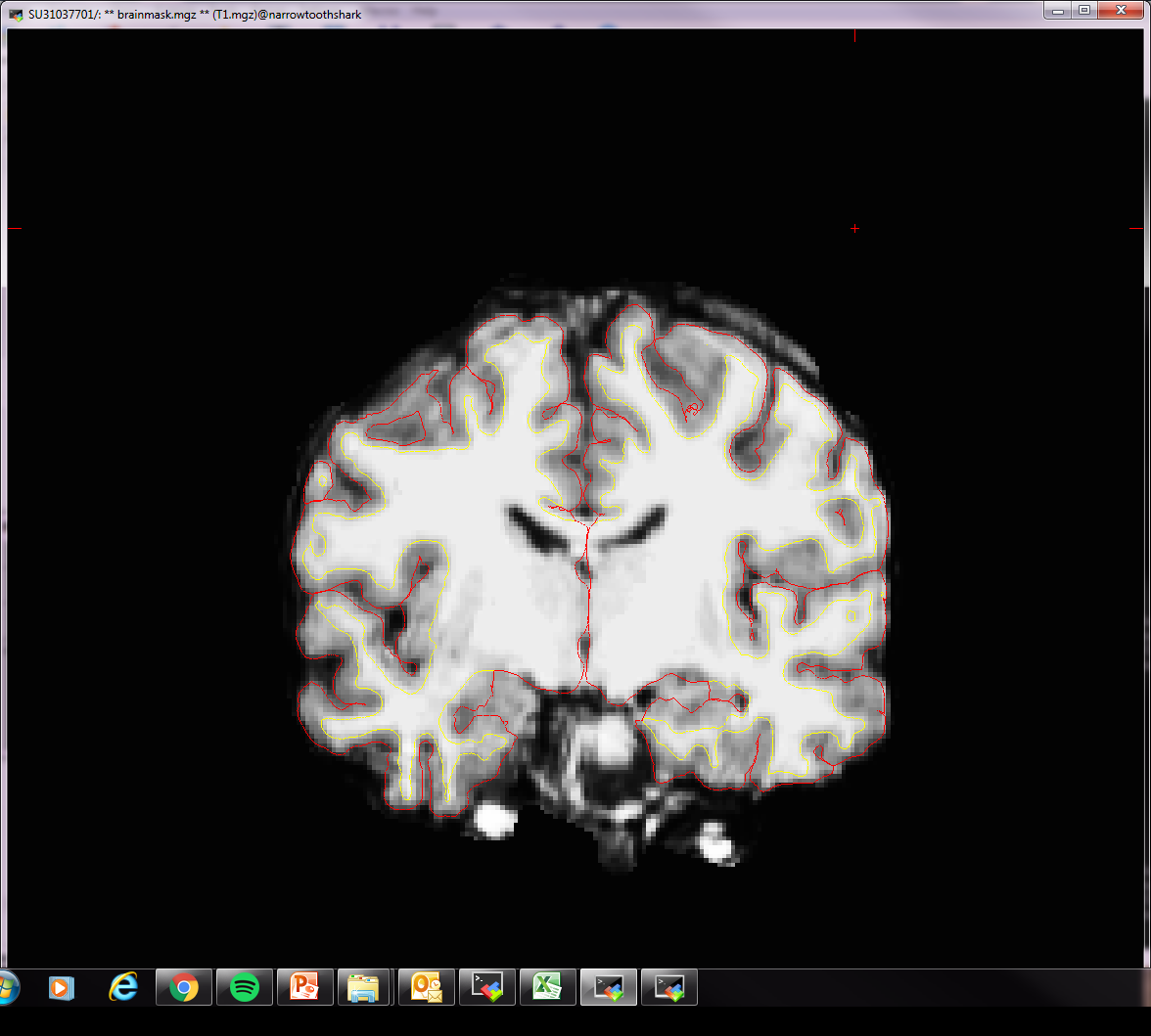

**Figure M4.** Example of a FreeSurfer processed image affected by movement.

Criterion 2: Is (part of) the temporal pole missing in the reconstruction?

*(0=NO, 1=YES)*

A particularly often-detected error is missing pial surface reconstruction at the temporal poles. An example can be seen in Figure M5. The pial surface of the temporal pole should be constructed from the first visible slice of the temporal pole. Although in Figure M5 the left temporal pole is clearly visible, the pial surface (red line) is reconstructed (highlighted by left square). If this is visible on three or more consecutive slices *Temp Pole Miss LH* (column D) is scored “1”. The right temporal pole showed poor reconstructed (highlighted by right square), however this is not observed on three consecutive slices, so *Temp Pole Miss RH* (column E) is scored as “0”.

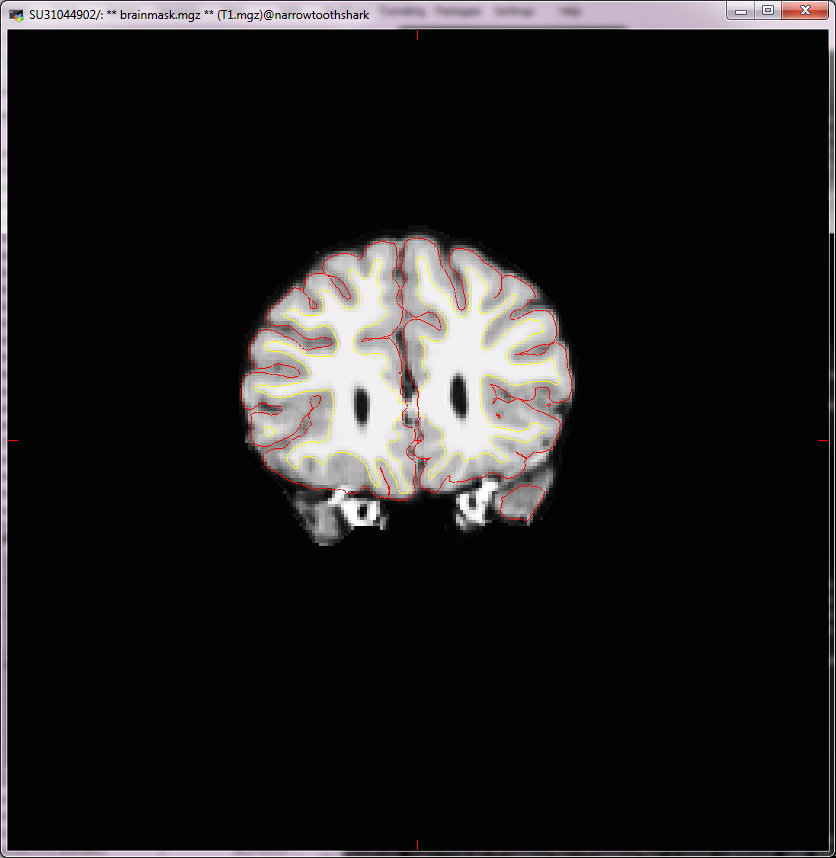

**Figure M5.** Example of temporal pole missing in reconstruction.

Criterion 3: Is non-brain tissue (e.g. dura/skull) included in the reconstruction of pial surface (red line)?

*(0=NO, 1=YES)*

Some brainmask.mgz files show skull or dura (see example in Figure M6). If dura is included in the reconstruction of the pial surface (see example on the right), *Dura LH* (column F) is scored “1”. If no dura is included in the reconstruction of the pial surface (see example on the left), *Dura RH* (column G) is scored “0”.

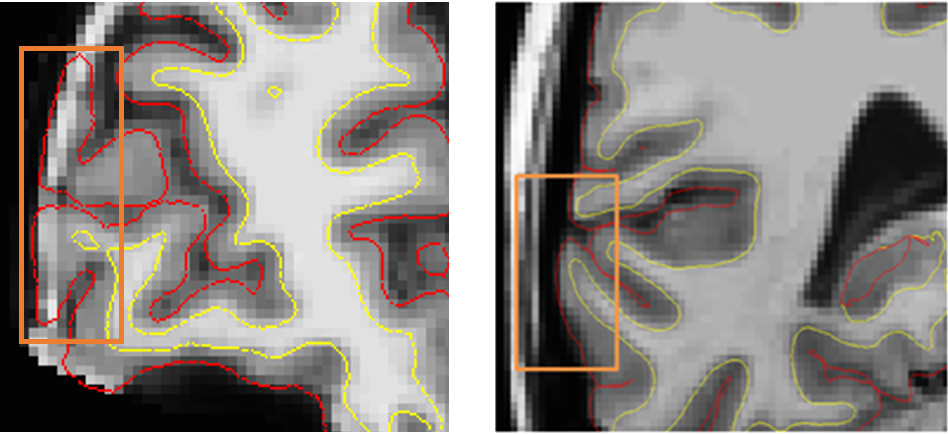

**Figure M6.** Example of dura included (left) and not included (right) in pial surface reconstruction.

Criterion 4: Are parts of the cortex missing in the reconstruction (other than temporal poles)?

*(0=NO, 1=YES)*

This criterion is aimed to evaluate whether parts of the cortex were not included in the reconstruction. We distinguished between anterior, superior/medial, and posterior areas of the brain. Again, this criterion is scored separately for the left and right hemispheres and is only indicated with a “1” if reconstruction errors are observed on 3 consecutive slices. An example of a slice with poor reconstruction is provided in Figure M7. The pial surface (indicated by the red line) should follow the outer border of the cortex as close as possible. The pial surface (red line) in the left hemisphere in Figure M7 did not follow the outer border of the cortex, leaving out some folds of the cortex. Therefore, *Missing Superior/Medial LH* (column J) has to be scored “1”. In comparison, the right hemisphere is reconstructed well, and *Missing Superior/Medial RH* (column K) has to be scored “0”.

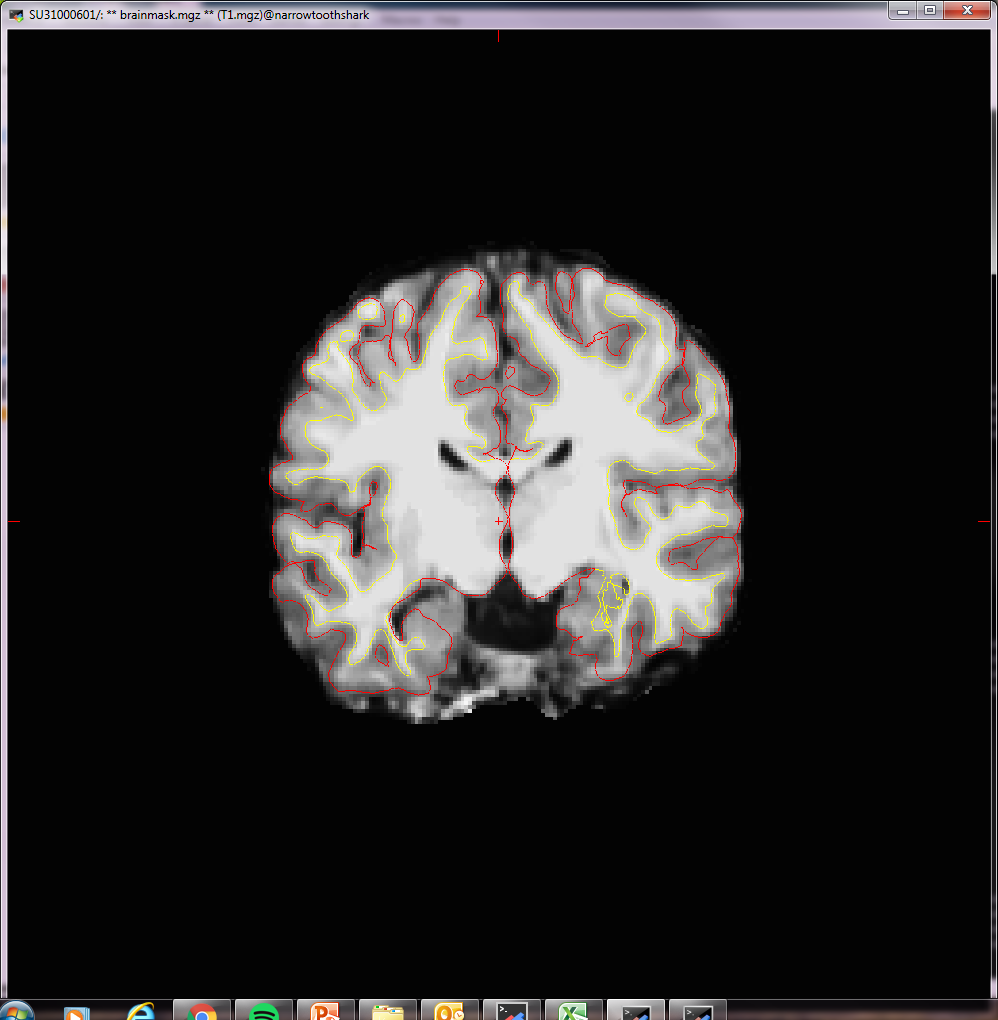

**Figure M7.** Example of cortical reconstruction error missing cortical brain tissue.

**2.4 Final score**

Each scan is rated with a final score, based on the assessment criteria described above. Final scores range from 1-4 and gives information on the quality of the scan:

1 = Excellent

2 = Good

3 = Doubtful

4 = Failed

In general, if all criteria were rated with 0, the scan should get a final score of 1. More “1” / Yes scores in the criteria lead to adjustment of the final score to Good, Doubtful or Failed. For example, a final score of 2 (Good) should be given if there were minor parts of the cortex not included in the reconstruction (e.g. missing anterior). If an additional criterion would also be violated (e.g. temporal pole or superior/medial missing, dura included) or if major parts are missed in the reconstruction, we recommend scoring the scan as a 3 (Doubtful). If there were many problems in multiple criteria (regardless of motion) we rated the scan as 4 (Failed). However, note that it is impossible to determine exact cutoffs between categories so they are sometimes arbitrary (stressing the added value of comparing manual scores with automated Qoala-T ratings). If the criterion *Excessive movement* is scored with 1 (i.e. there is excessive movement), we would recommend that the scan is always scored as 4 (Failed).
